# Topological regulation of the estrogen transcriptional response by ZATT-mediated inhibition of TOP2B activity

**DOI:** 10.1101/2024.01.22.576640

**Authors:** José Terrón-Bautista, María del Mar Martínez-Sánchez, Laura López-Hernández, Ananda Ayyappan Vadusevan, Mario García-Domínguez, R. Scott Williams, Andrés Aguilera, Gonzalo Millán-Zambrano, Felipe Cortés-Ledesma

**Affiliations:** Centro Andaluz de Biología Molecular y Medicina Regenerativa-CABIMER, Universidad de Sevilla-CSIC-Universidad Pablo de Olavide, Sevilla, Spain; Genome Integrity and Structural Biology Laboratory, National Institute of Environmental Health Sciences, National Institutes of Health, US Department of Health and Human Services, Research Triangle Park, NC 27709, USA; Departamento de Genética, Universidad de Sevilla, Spain; Topology and DNA Breaks Group, Spanish National Cancer Centre (CNIO), Madrid 28029, Spain

**Author notes:** Correspondence (G.M.-Z), (F.C.-L.).

## Abstract

Human type-II topoisomerases, TOP2A and TOP2B, remove transcription associated DNA supercoiling, thereby affecting gene-expression programs, and have recently been associated with 3D genome architecture. Here, we study the regulatory roles of TOP2 paralogs in response to estrogen, which triggers an acute transcriptional induction that involves rewiring of genome organization. We find that, whereas TOP2A facilitates transcription, as expected for a topoisomerase, TOP2B limits the estrogen response. Consistent with this, TOP2B activity is locally downregulated upon estrogen treatment to favor the establishment and stabilization of regulatory chromatin contacts, likely through an accumulation of DNA supercoiling. We show that estrogen-mediated inhibition of TOP2B requires estrogen receptor α (ERα), a non-catalytic function of TOP2A, and the action of the atypical SUMO-ligase ZATT. This mechanism of topological transcriptional-control, which may be shared by additional gene-expression circuits, highlights the relevance of DNA topoisomerases as central actors of genome dynamics.

## INTRODUCTION

Around 75% of breast cancers are driven by the estrogen receptor α (ERα)-dependent transcriptional response. The activity of ERα largely depends upon the presence of estrogen hormones, which mediate its homodimerization and subsequent binding to more than 10,000 different sites across the genome.^1,2^ ERα-binding sites (ERBSs) are predominantly located at enhancers, whose transcription is also regulated in response to estrogen.^2–4^ Whereas the exact function of estrogen-regulated enhancer RNAs (eRNAs) themselves remains controversial, it seems clear that enhancer and target-gene transcription kinetics strongly correlate.^3–5^ In line with this, it has become apparent that the transcriptional response to estrogen functions by extensive ERα-mediated chromatin looping that brings enhancers and target genes together for coordinated regulation. ^6–9^ The association between dysregulated ERα-mediated long-range interactions and endocrine therapy resistance in breast cancer underscores their physiological relevance.^10,11^

Chromatin contacts are mediated, or at least stimulated, by cohesin, a ring-shaped complex that, by holding DNA segments together, establishes genome organization into loops.^12,13^ This is a dynamic process in which, from its loading site, cohesin initiates and gradually enlarges the chromatin loop (“loop extrusion”) until it becomes arrested at specific boundaries. Thus, contacts between functional elements can be regulated by cohesin loading sites and the position of barriers or hurdles that limit its extrusion. Interestingly, it has been reported that enhancers can recruit cohesin,^14,15^ while the presence of RNA polymerase II (RNA Pol II), both at enhancers and promoters, seems to counteract the loop extrusion process ^16,17^, thereby providing an interesting framework for the regulation of enhancer-promoter interactions.

Transcription-derived DNA supercoiling has also been proposed to favor regulatory interactions. This could occur simply by facilitating the physical proximity between the DNA elements, as shown both *in vitro* and using molecular simulations,^18,19^ but, interestingly, also by “pushing” the cohesin ring thereby promoting loop extrusion. ^20,21^ In mammalian cells, transcription-dependent DNA supercoiling is mainly relieved by the action of DNA topoisomerases I (TOP1) and II (TOP2A and TOP2B), through controlled cut-and-reseal mechanisms in which the enzyme covalently associates to DNA in a cleavage-complex intermediate.^22^ TOP1 physically interacts with RNA Pol II, but only becomes fully activated during transcription elongation, enabling local removal of DNA supercoiling directly associated with ongoing transcription.^23^ Whereas TOP2A and TOP2B have similar catalytic and structural properties, they are not functionally redundant and display specific expression patterns and different, yet partially overlapping, functions. ^22,24^ TOP2B is expressed throughout the cell cycle and has been mainly associated with transcription and, more recently, with genome architecture.^25–28^ In contrast, TOP2A is highly expressed during mitosis and has been traditionally linked to chromosome segregation, although this general view is challenged by recent reports suggesting additional relevant functions during transcription.^29,30^ Consistently, both TOP2A and TOP2B are enriched at transcriptional regulatory elements, such as enhancers and promoters.^31,32^

Accumulating evidence suggest that TOP2 paralogues play important roles during the induction of stimuli-responsive genes.^30,33–37^ However, their mechanism of action is still controversial and not completely understood. Here, we have systematically addressed the roles of TOP2A and TOP2B during the ERα-dependent transcriptional response in breast cancer MCF7 cells. Unexpectedly, we found that TOP2 paralogs have opposing functions in estrogen-mediated transcriptional regulation. Our results indicate that TOP2A, whose activity is upregulated at estrogen-induced genes, facilitates the estrogen response. This is in agreement with the expected function of a topoisomerase in relieving the topological stress derived from increased transcription. In contrast, we found that TOP2B limits the estrogen response, and that its activity is strikingly downregulated at ERα target regions upon estrogen stimulation. This repressive function of TOP2B seems to operate at the level of preventing enhancer-promoter contacts, likely through the removal of transcription-induced negative (-) DNA supercoiling, and is directly controlled by the TOP2-interacting SUMO ligase ZATT/ZNF451. Thus, we propose a novel mechanism of topological regulation by which ZATT-mediated inhibition of TOP2B allows the accumulation of (-) DNA supercoiling to favor regulatory chromatin contacts and full activation of the estrogen-dependent transcriptional response.

## RESULTS

### Human TOP2 paralogs play different roles during the estrogen transcriptional response

To examine functions of TOP2A and TOP2B during the estrogen-dependent transcriptional response, we transfected hormone-depleted MCF-7 cells with paralog-specific small interfering RNAs (siRNA) (Figure S1A). We then treated cells with 17β-estradiol (E2) for 45 minutes and performed RNA Pol II chromatin immunoprecipitation experiments followed by high-throughput DNA sequencing (ChIPseq). Of note, all ChIPseq experiments included in this study rely on a mouse chromatin spike-in control, which allows for quantitative comparisons between conditions and can reveal uniform changes in occupancy.^38^ On a global scale, we found 359 protein-coding genes to display induced RNA Pol II occupancy (log2FC > 1; p-value < 0.05) after 45 minutes of E2 treatment when compared to untreated conditions (Figure S1B). These included classical estrogen-regulated genes, such as *GREB1* and *RET*, validating the use of RNA Pol II recruitment as an indicator of estrogen-mediated gene induction, and confirming that this short E2 treatment allows for the detection of direct and immediately responding target genes. We also detected 264 enhancers (as identified in^3^) to be induced upon estrogen treatment (Figure S1C). We selected these sets of E2-induced genes and enhancers, and, for comparison purposes, random sets containing the same number of non-responsive elements.

As can be seen in Figure 1A, metaplot and quantification analyses showed a clear estrogen-dependent increase of RNA Pol II occupancy at both E2-induced genes and enhancers. TOP2A depletion, however, specifically limited E2-induced RNA Pol II recruitment (Figure 1A and 1B), in agreement with a positive role of TOP2A in mediating the estrogen response. In contrast, depleting TOP2B did not decrease estrogen-mediated RNA Pol II recruitment along estrogen-induced genes or enhancers (Figure 1C and 1D). As a matter of fact, we observed significantly increased accumulation specifically at the promoters (Figure 1E). These results, contrary to a previous report assigning a positive function of TOP2B in triggering the estrogen response^33^, suggest that TOP2B may limit estrogen signaling, at least at the level of RNA Pol II recruitment to responsive promoters.

**Figure 1.**
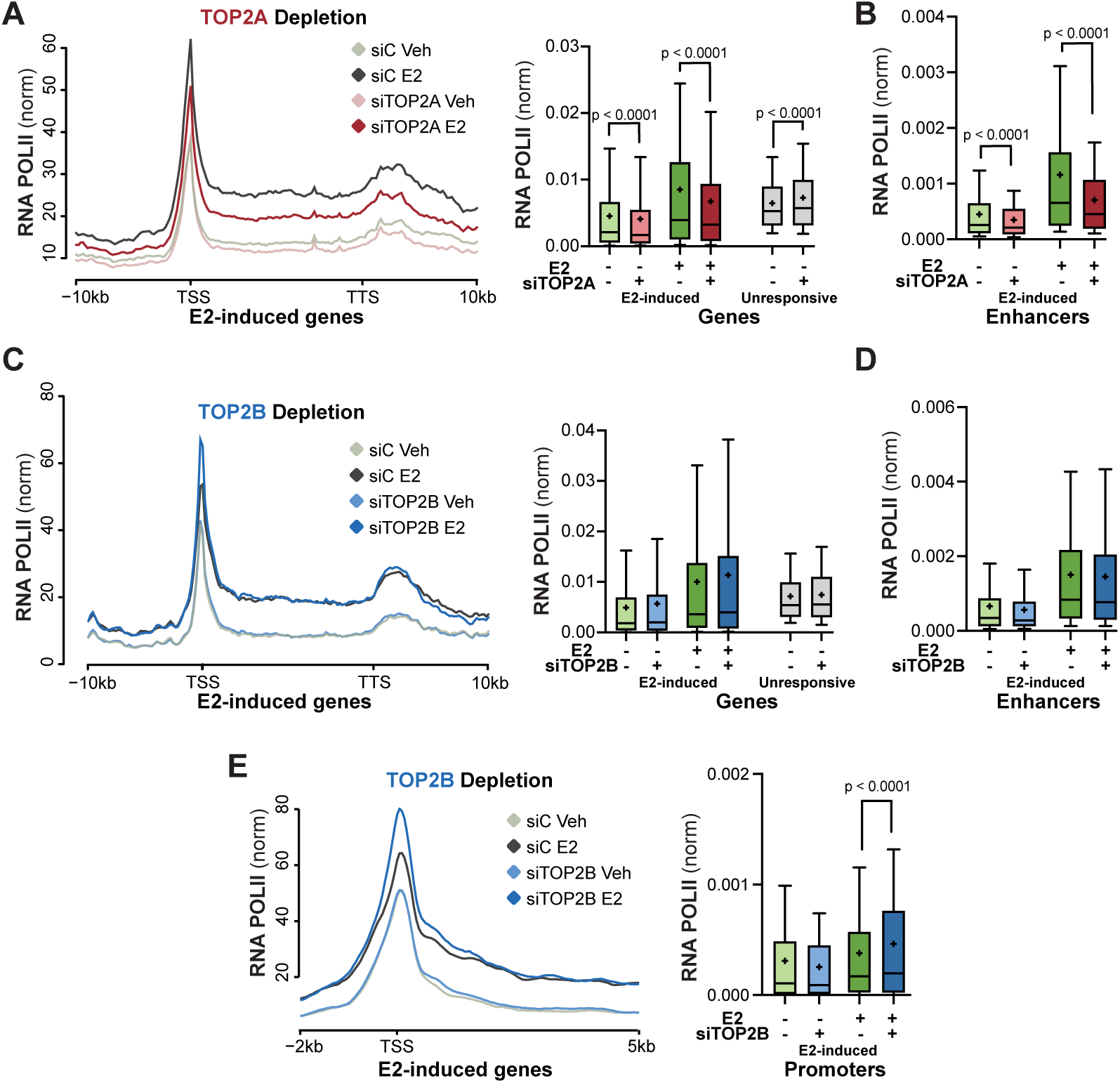
Human TOP2 paralogs play opposite roles during the estrogen transcriptional response. (A) Metaplot (left) and quantification (right) of average RNA Pol II ChIPseq enrichment, represented as normalized reads per genome coverage (RPGC), along E2-responsive genes in hormone-depleted MCF7 cells transfected with non-targeting control (siC) or TOP2A-specific (siTOP2A) siRNAs in untreated control conditions (vehicle, Veh) or upon E2 treatment (10 nM, 45 min). Quantification of average RNA Pol II ChIPseq enrichment, represented as normalized reads per genome coverage (RPGC) at shown E2-responsive genes. (B) Quantification of average RNA Pol II ChIPseq enrichment, represented as normalized reads per genome coverage (RPGC) at E2-responsive enhancers. Other details as in “A”. (C) As in “A” in cells transfected with non-targeting control (siC) or TOP2B-specific (siTOP2B) siRNAs. (D) As in “B” in cells transfected with non-targeting control (siC) or TOP2B-specific (siTOP2B) siRNAs. (E) As in “C” at oriented TSS of E2-responsive genes. Quantification at shown TSS (- 100 bp/ +400 bp). Statistical analysis was performed using multiple Wilcoxon test with Holm-Sídák correction (adjusted p value is shown).

To validate these surprising findings, and to address whether the effects observed were dependent on the activity of the TOP2 paralogs, we used catalytic inhibitors such as merbarone and ICRF-187, which have been reported to display partial differential inhibitory potencies towards TOP2A and TOP2B. ^30,39^ Indeed, ICE (*In vivo* Complex of Enzyme) assays, which measure the accumulation of TOP2 cleavage complexes (TOP2cc) upon a brief treatment with the TOP2cc stabilizer etoposide as a proxy of TOP2 activity, showed a preference of merbarone and ICRF187 for inhibiting TOP2A and TOP2B, respectively (Figure S1D). We then used these experimental conditions of preferential TOP2-paralog inhibition and monitored changes in E2-induced RNA Pol II recruitment. As a matter of fact, we found that merbarone treatment also resulted in reduced E2-mediated RNA Pol II recruitment to estrogen-induced genes and enhancers (Figure S1E and S1F), confirming the results obtained upon TOP2A depletion. In contrast, ICRF-187 treatment caused increased RNA Pol II recruitment in response to E2 treatment, in this case not only at the promoters, but also along estrogen-induced genes and enhancers (Figure S1G and S1H). Thus, in the case of acute TOP2B inhibition, the increased RNA pol II recruitment observed upon long-term depletion is further translated into higher transcriptional levels. Collectively, the above findings suggest that the two TOP2 paralogs play different roles during the estrogen response, with TOP2A facilitating transcription and TOP2B restricting at least RNA pol II recruitment to promoter regions.

### TOP2B activity removes (-) DNA supercoiling and restrains ERα-mediated contacts between regulatory regions

While the results after TOP2A depletion and inhibition are consistent with the expected role of topoisomerases as facilitators of transcription, the effects observed upon TOP2B targeting are suggestive of an additional uncharacterized function, which we aimed at investigating in more detail. Given the classical role of TOP2B in the removal of topological stress associated with transcription,^24^ we decided to measure how TOP2B inactivation affected the accumulation of DNA supercoiling in our experimental system. We used biotin-psoralen (BP) photobinding, which preferentially intercalates into negatively (-) supercoiled DNA,^40,41^ and subsequent detection and quantification in individual nuclei with fluorescently-labeled streptavidin. In agreement with transcription being a major source of (-) DNA supercoiling in cells,^42,43^ we found that BP incorporation was strongly reduced upon preincubation with the global transcriptional inhibitor triptolide (Figure 2A). In contrast, short-term inhibition of TOP2B activity by ICRF187 treatment caused a significant increase in BP incorporation (Figure 2A), indicative of a major function of TOP2B in removing (-) DNA supercoiling. Importantly, no changes in chromatin accessibility, which can also affect BP incorporation, were observed upon ICRF-187 treatment, neither at responsive regions nor globally (Figure S2A and S2B). Finally, a combination with ERα immunofluorescence analysis revealed that ICRF187 treatment also increased (-) DNA supercoiling levels specifically at nuclear regions with strong ERα staining, which were in fact particularly enriched in BP photobinding when compared to the rest of the nucleus (Figure 2B), suggesting that ERα signaling constitutes a local context that is prone to the accumulation of (-) DNA supercoiling. These results link TOP2B activity to the removal of transcription-induced (-) supercoiling, and suggest that this may be particularly relevant in the context of ERα signaling.

**Figure 2.**
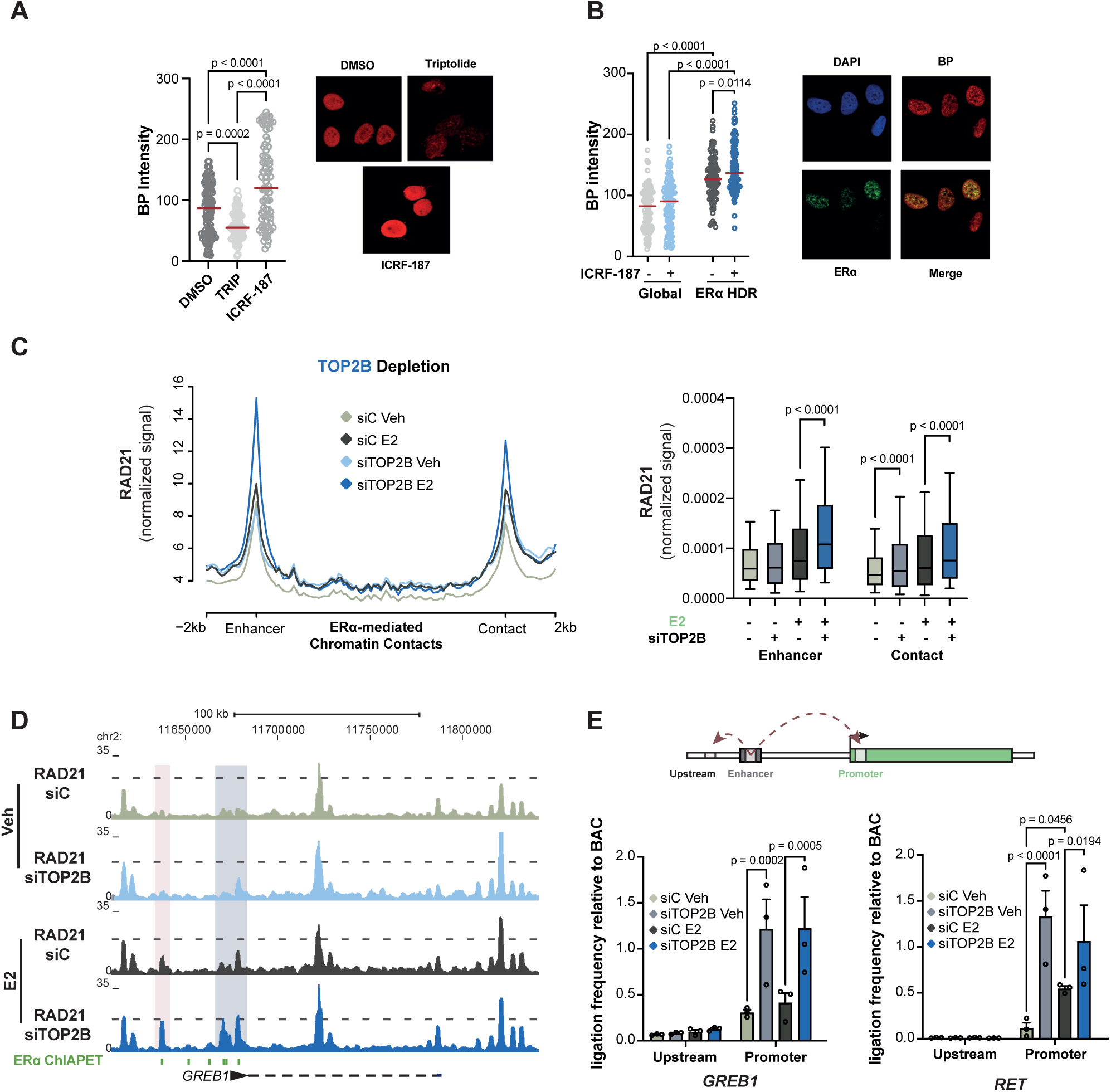
TOP2B activity removes (-) DNA supercoiling and restrains ERα-mediated contacts between regulatory regions. (A) Immunofluorescence analysis of BP photobinding after triptolide (10 uM, 5 h) or ICRF-187 (100 uM, 30 min) treatments. The distribution and median (red bar) of BP intensity in individual nuclei from 3 independent experiments (left) and a representative image (right) are shown. Statistical analysis was performed using One-way ANOVA. (B) Immunofluorescence analysis of BP photobinding and ERα with and without ICRF-187 treatment (30 min, 100 uM), as indicated. The distribution and median (red bar) of BP intensity in entire individual nuclei (Global) and specifically at regions of high ERα density (ERα HDR) from 3 independent experiments (left) and a representative image (right) are shown. (C) Metaplot (left) and quantification (right) of RAD21 ChIPseq normalized counts (RPGC) at ERα-mediated enhancer chromatin contacts (+/- 500 base pairs) in hormone-depleted MCF7 cells transfected with non-targeting control (siC) or TOP2A-specific (siTOP2A) siRNAs in untreated control conditions (vehicle, Veh) or upon E2 treatment (10 nM, 45 min). Statistical analysis was performed using multiple Wilcoxon test with Holm-Sídák correction (adjusted p value is shown). (D) Genome browser view at the *GREB1* regulatory landscape showing RAD21 ChIPseq normalized signal from datasets described in “C”. Enhancer (as determined by previous ERα ChIAPET experiments ^6^) and promoter of *GREB1* are indicated by red and grey boxes, respectively. (E) 3C qPCR experiments showing enhancer-promoter contact frequency at the *GREB1* (left) and *RET* (right) loci in hormone-depleted MCF7 cells transfected with non-targeting control (siC) or TOP2B-specific (siTOP2B) siRNAs in untreated control conditions (vehicle, Veh) or upon E2 treatment (10 nM, 45 min). A general scheme of the contacts analyzed is shown (top). Signal is normalized to the intrinsic ligation frequency of the same sequences cloned in BAC plasmids. Mean + SEM of three independent experiments is shown. Statistical analysis was performed using One-way ANOVA.

DNA supercoiling has been proposed to favor distal chromatin contacts, either directly through the formation of superhelical structures,^18,19^ or indirectly by driving cohesin movement.^20,21^ Furthermore, TOP2B binding is tightly linked to 3D genome organization,^26–28^ although its specific role within this context is still poorly understood. Based on this, we decided to analyze whether TOP2B depletion may affect the regulatory chromatin contacts taking place in response to estrogen. To do so, we first assessed the distribution of the cohesin complex by performing RAD21 ChIPseq experiments upon E2 treatment. We focused our analysis on estrogen-induced ERα-mediated chromatin contacts of enhancer regions, as previously identified by ERα ChIA-PET experiments.^6^ As reported before,^44,45^ we found that E2 treatment globally increased RAD21 binding at E2-mediated enhancer-contact pairs (Figure 2C), which can be clearly observed at the *GREB1* locus as an illustrative example (Figure 2D). Strikingly, we observed that TOP2B depletion further increased estrogen-induced cohesin binding at these regions (Figure 2C and 2D). Furthermore, TOP2B depletion, on its own, resulted in an increased cohesin occupancy at the enhancer contact point, but not at the enhancer itself, even under non-induced conditions (Figure 2C). We conclude that TOP2B negatively affects cohesin occupancy at ERα regulatory contact regions.

To test whether the effect of TOP2B depletion on cohesin distribution reflected actual changes in 3D chromatin organization, we performed chromatin conformation capture (3C) qPCR experiments and analyzed the well characterized ERα-mediated enhancer-promoter chromatin contacts taking place at the *GREB1* and *RET* loci.^6,8^ We found that these enhancer-promoter interactions were, as expected, increased by E2 stimulation, but also by TOP2B depletion on its own, irrespective of estrogen treatment (Figure 2E). Collectively, the above findings suggest that TOP2B function restrains cohesin-mediated regulatory chromatin contacts taking place during the estrogen response, and are consistent with models in which this could be exerted by controlling negative DNA supercoiling.^20,21^ TOP2B depletion/inhibition would thus generate a topological environment that facilitates regulatory chromatin interactions, providing a molecular explanation for its repressive functions in the estrogen-dependent transcriptional response.

### TOP2B is locally inactivated during the estrogen response

We have uncovered that while TOP2A facilitates estrogen-mediated RNA Pol II recruitment, TOP2B limits it, likely by reducing long-range chromatin contacts. Therefore, we wondered whether these functions could be exploited by the cell to physiologically control the estrogen response. To address this, we first monitored genome-wide changes in TOP2A and TOP2B binding upon E2 treatment by performing ChIP-seq experiments. In agreement with previous studies^30–32^, we found that both TOP2A and TOP2B were enriched at enhancers and promoters (Figure S3A), while TOP2B was additionally found at insulator regions. Moreover, both of their occupancies strongly correlated with RNA Pol II density at transcribed regions (Figure S3B), which is consistent with their role in the resolution of topological stress derived from transcription.^24^ When focusing on the estrogen response, we found that TOP2A was strongly recruited to E2-induced genes and enhancers upon estrogen treatment (Figure 3A), and this was indeed specific to responsive regions (Figure 3B). In contrast, we observed that TOP2B binding remained practically unaltered at these regions after E2 stimulation (Figure 3C), although there was a slight but significant redistribution form non-response to responsive genes (Figure 3D). Note that the specificity of the antibodies was confirmed by ChIP-qPCR analysis at the *GREB1* enhancer and promoter regions (Figure S3C).

**Figure 3.**
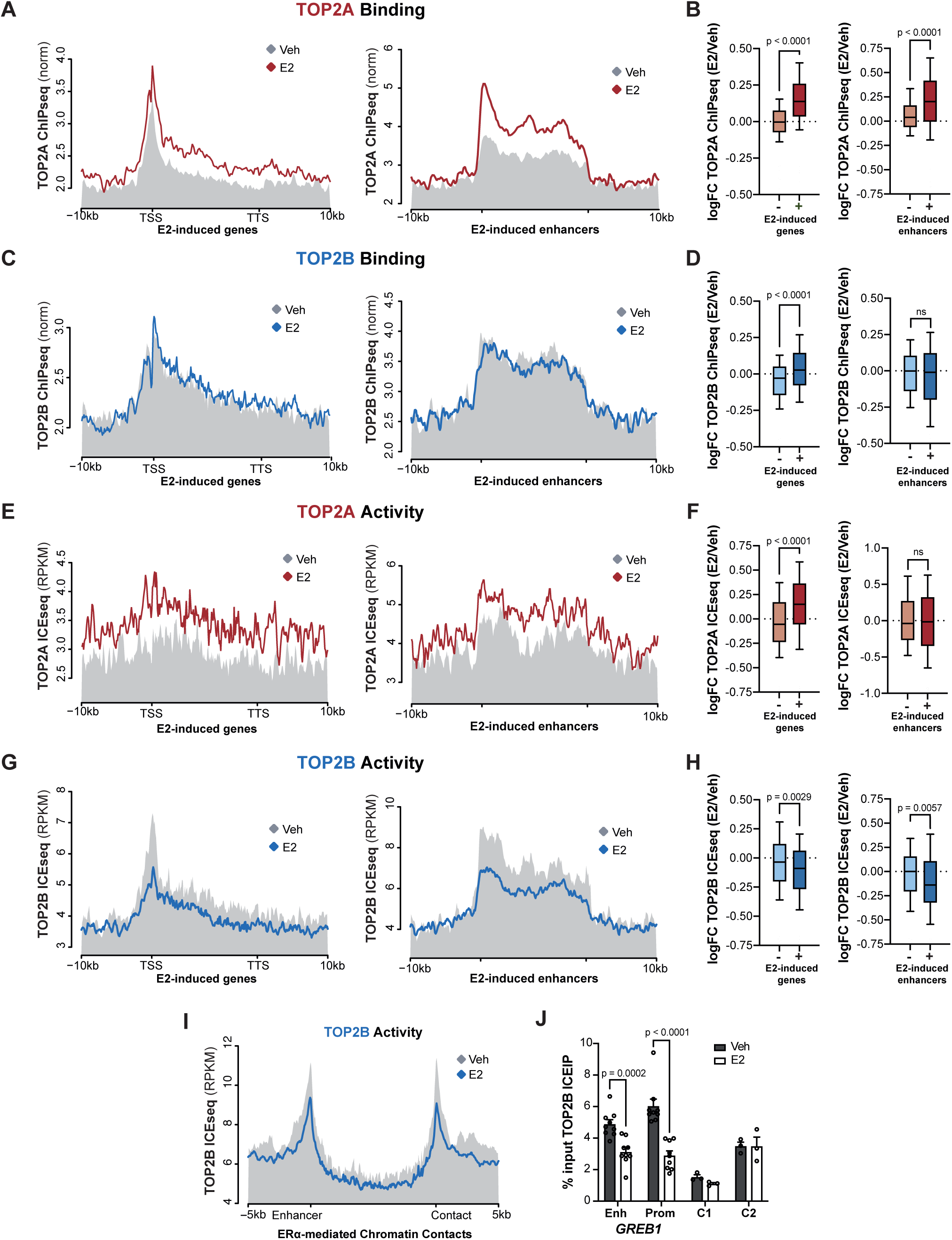
TOP2B is locally inactivated during the estrogen response. (A) Metaplot depicting average TOP2A ChIPseq enrichment (RPGC) along E2-responsive genes (left) and enhancers (right) under control (vehicle, Veh) conditions and upon E2 treatment (45 min, 10 nM). (B) Quantification of the fold change of TOP2A ChIPseq signal between E2 and vehicle conditions at E2-responsive and non-responsive genes (left) and enhancers (right). Boxplots represents 10-90 percentile and median of the signal. Statistical analysis was performed using Wilcoxon test. (C) As in “A” for TOP2B ChIPseq signal. (D) As in “B” for TOP2B ChIPseq signal. (E) As in “A” for TOP2A ICEseq signal. (F) As in “B” for TOP2A ICEseq signal. (G) As in “A” for TOP2B ICEseq signal. (H) As in “B” for TOP2B ICEseq signal. (I) Metaplot representation of TOP2B ICEseq distribution (RPKM) at ERα-mediated enhancer chromatin contacts, as determined by previous ERα ChIAPET experiments.^6^ Hormone-depleted cells were treated with 10 nM E2 for 45 minutes. The regions between the contact points +/- 5 kilobases are shown. (J) ICEIP qPCR experiments showing TOP2B activity at the indicated locations of the *GREB1* regulatory landscape. Hormone-depleted cells were treated with 10 nM E2 for 45 minutes. Control sites are nearby regions characterized by TOP2B binding but not ERα recruitment. Data are represented as mean + SEM of at least three independent experiments. Statistical analysis was performed using Two-way ANOVA.

We next considered the possibility that the function of TOP2 paralogs, and of TOP2B in particular, was further regulated at the level of its catalytic activity. To address this, we first performed traditional ICE assays upon a brief incubation with etoposide, which, as described above, can be taken as a proxy of TOP2 activity. However, we did not find significant differences in neither TOP2Acc nor TOP2Bcc accumulation upon E2 stimulation (Figure S3D), ruling out global changes in TOP2 activity during the estrogen response. Then, to investigate local effects, we used an adapted version of the ICE protocol that allows for paralog-specific TOP2cc immunoprecipitation and tagmentation-based DNA library preparation (ICE-seq; see STAR methods). Specificity of this new assay was confirmed by performing ICE qPCR experiments at the *GREB1* enhancer and promoter regions upon paralog-specific siRNA-mediated knockdown (Figure S3E). Global distribution of the ICE-seq signal showed, in general, a significant consistency with ChIP-seq data for both TOP2A and TOP2B with a strong accumulation at regulatory regions (Figure S3A). However, we also found relevant differences that likely reflect specific modulation of their activities. For example, peaks of TOP2B activity were increased at promoters and reduced at insulator regions. Regarding the estrogen response, metaplot representation of ICE-seq signal upon E2 treatment showed that TOP2A activity increased both at estrogen-induced genes and enhancers (Figure 3E). Quantification and comparison with non-responsive elements, however, showed that only responsive genes displayed a specific increase in TOP2A activity (Figure 3F), despite it being similarly recruited to both genes and enhancers (Figure 3B). We conclude that the estrogen response is accompanied by a local increase in the activity of TOP2A, likely dealing with the topological stress derived from induced transcription.

In contrast to this, we found that TOP2B ICE-seq signal showed a marked reduction upon E2 treatment both at estrogen-induced genes and enhancers (Figure 3G), and this reduction was significant when compared to non-responsive elements (Figure 3H). Furthermore, we also observed a reduction at ERα-mediated enhancer contacts (Figure 3I). The decrease in TOP2B ICE signal was confirmed by qPCR analysis at the enhancer and promoter of the *GREB1* gene (Figure 3J; see also genome-browser view in Figure S3F), was still observed in the presence of flavopiridol and triptolide transcriptional inhibitors (Figure S3G), and did not reflect increased TOP2Bcc degradation, since TOP2B ChIPseq signal was not affected by the brief etoposide treatment used (Figure S3H). We therefore conclude that the estrogen response is accompanied by a downregulation of TOP2B activity at E2 responsive elements, including those that mediate regulatory chromatin contacts. It is worth noting that this decrease in TOP2B activity occurred in the context of high topological stress, as evidenced by the strong increase in TOP1 activity observed in responsive regions upon estrogen treatment (Figure S3I). Collectively, these results indicate that the activity of TOP2 paralogs is differentially regulated during the estrogen response, with an increase in TOP2A and a decrease in TOP2B activities that are in agreement with their respective stimulatory and repressive functions. Most importantly, the downregulation of TOP2B activity suggests a novel mechanism of transcriptional regulation that could operate by controlling regulatory chromatin contacts.

### Downregulation of TOP2B activity depends on ERα and TOP2A

We next sought to uncover the molecular mechanism(s) by which the activity of TOP2B was downregulated in response to estrogen. Given that this downregulation specifically occurred at estrogen-induced loci, we wondered whether TOP2B inactivation required ERα recruitment. As mentioned above, ERα-binding sites are predominantly located at enhancers,^2^ so virtually all estrogen-induced enhancers show ERα recruitment under our experimental conditions. However, when classifying estrogen-induced protein coding genes depending on whether or not their promoters displayed ERα recruitment, we found that TOP2B activity was specifically downregulated at those bound by ERα (Figure 4A). Consistent with this, and as an illustrative example, TOP2B ICE-IP qPCR analysis at the estrogen-responsive gene *P2RY2* revealed that TOP2B activity was downregulated only at the enhancer, which shows ERα binding, but not at the promoter, which does not (Figure 4B). In contrast, both the enhancer and promoter regions of the *GREB1* locus, which display strong ERα recruitment, showed downregulation of TOP2B activity (Figure 4B). To establish cause-effect relationships, we next treated cells with fulvestrant, a competitive ERα antagonist that triggers its degradation,^46^ and analyzed TOP2B activity regulation in response to estrogen. Despite incomplete ERα degradation (Figure S4A), we observed that fulvestrant pre-treatment partially suppressed E2-induced TOP2B downregulation (Figure 4C), further supporting the notion that ERα recruitment is important for TOP2B inactivation.

**Figure 4.**
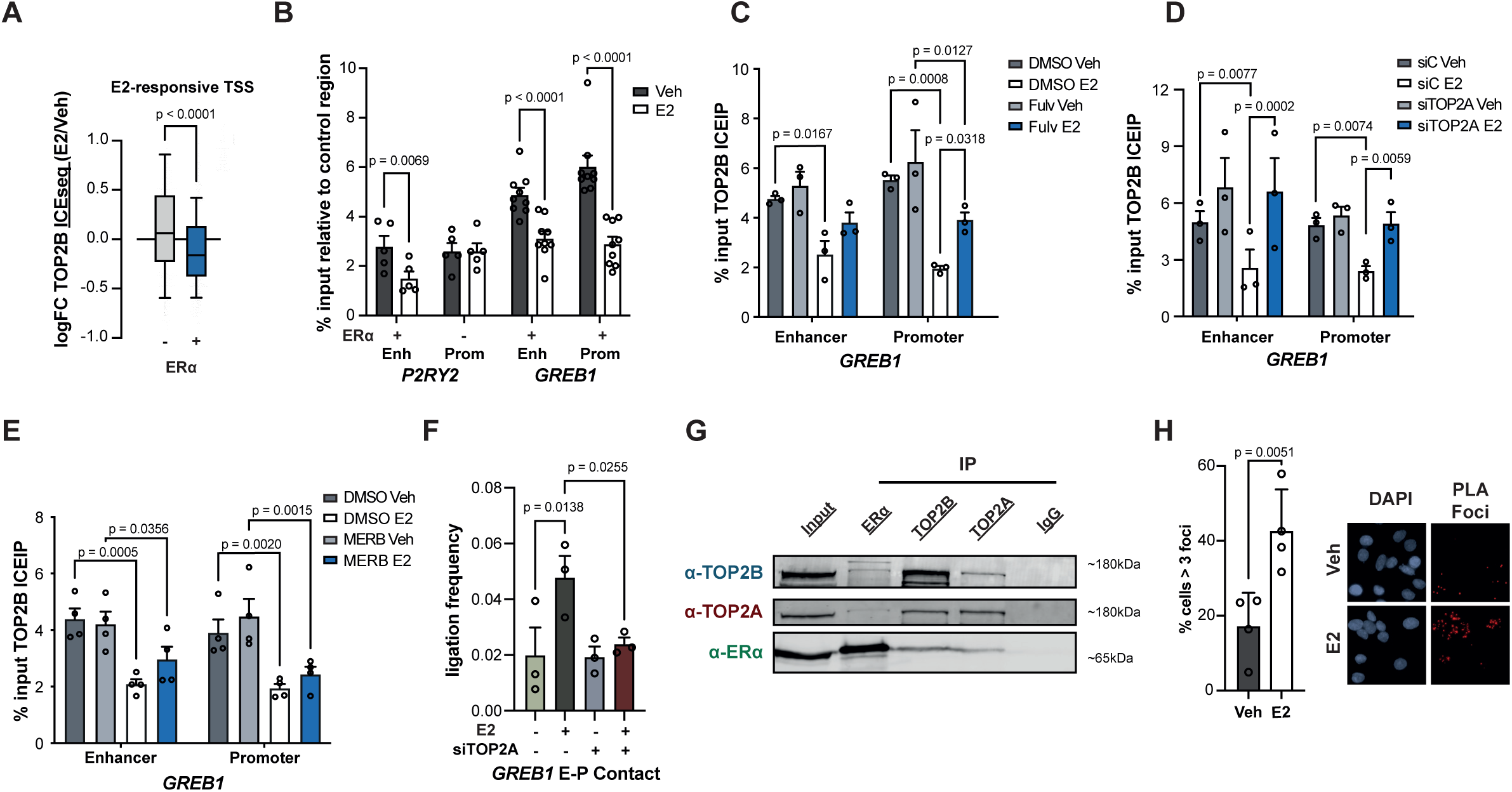
Downregulation of TOP2B activity depends on ERα and TOP2A. (A) Average fold change (median of log2) of TOP2B ICEseq signal (RPKM) at E2 responsive transcription start sites (TSS) (+/- 5 kb) upon E2 treatment (45 min, 10 nM). TSS were classified into non-ERα-bound or ERα-bound (logFC > 1.5). Boxplots represents 10-90 percentile and median of the signal. Statistical analysis was performed using Wilcoxon test. (B) ICEIP qPCR experiments showing TOP2B activity at different regions of *P2RY2* and *GREB1* loci, which recruit ERα or not as indicated, in hormone-depleted MCF7 cells in untreated control conditions (vehicle, Veh) or upon E2 treatment (10 nM, 45 min). Mean + SEM of at least three independent experiments is shown. Statistical analysis was performed using Two-way ANOVA. (C) As in (B) in enhancer and promoter regions of *GREB1* in cells that were pre-treated (Fulv) or not (DMSO) with fulvestrant (100 nM, 3 h). (D) As in (C) cells transfected with non-targeting control (siC) or TOP2A-specific (siTOP2A) siRNAs. (E) As in (C) in cells that pre-treated (Merb) or not (DMSO) with merbarone (100 nM, 90 min). (F) 3C qPCR experiments showing enhancer-promoter contact frequency at the *GREB1* locus in hormone-depleted MCF7 cells transfected with non-targeting control (siC) or TOP2A-specific (siTOP2A) siRNAs. Details as in Figure 2E. (G) Immunoblot of TOP2B (top), TOP2A (middle) and ERα (bottom) in extracts of MCF7 cells (Input) and immunoprecipitated material (IP) using anti-ERα, -TOP2B, - TOP2A and control IgG antibodies, as indicated. (H) Quantification (left) and representative image (right) of proximity ligation assays (PLA) using TOP2A and TOP2B specific antibodies in hormone-depleted MCF7 cells treated with vehicle (Veh) or E2 treated (10 nM, 45 min). Data are represented as Mean + SEM of the percentage of cells displaying more than 3 PLA foci in four independent experiments is shown. Statistical analysis was performed using Chi-square with Yates correction.

As described above, TOP2A is also recruited to estrogen-responsive regions upon E2 treatment. Interestingly, TOP2A recruitment to promoters strongly correlated with ERα recruitment, even more than with increases in RNA Pol II density (Figure S4B), suggesting that it does not merely depend on increased transcription. We thus wondered whether TOP2A was also involved in the estrogen-induced downregulation of TOP2B activity, perhaps due to some kind of crosstalk between the two paralogs. To test this, we performed TOP2B ICE-IP qPCR analysis at the *GREB1* locus upon siRNA-mediated knockdown of TOP2A, which, strikingly, completely suppressed downregulation of TOP2B activity in response to estrogen (Figure 4D). Interestingly, however, TOP2A inhibition by merbarone treatment did not significantly affect the estrogen-induced decrease in TOP2B activity (Figure 4E), arguing that TOP2B inactivation depends on TOP2A, but not on its catalytic activity. Furthermore, we found that TOP2A depletion completely suppressed the increase in the enhancer-promoter chromatin contact taking place at the *GREB1* locus upon E2 stimulation (Figure 4F). We conclude that recruitment of ERα to its responsive elements, together with a non-catalytic function of TOP2A, trigger a local downregulation of TOP2B activity that facilitates regulatory chromatin contacts.

### ZATT inhibits TOP2B activity to facilitate the estrogen response

To shed some light into the possible mechanism by which ERα and TOP2A inactivate TOP2B, we performed co-immunoprecipitation experiments. Pulling down endogenous ERα co-immunoprecipitated TOP2B, as previously reported,^33^ but also TOP2A (Figure 4G). Interaction between the three factors was, as a matter of fact, confirmed in reciprocal pull-down experiments (Figure 4G). These results, together with their respective ChIP-seq profiles at estrogen-responsive elements, suggest that ERα can simultaneously interact with TOP2A and TOP2B paralogs on chromatin. In fact, *in situ* proximity ligation assays (PLA) indicated an association between TOP2A and TOP2B, which was significantly increased upon E2 stimulation (Figure 4H). Interestingly, part of the fraction of TOP2B that was associated with ERα showed an upward mobility shift (Figure 4G), suggestive of possible post-translational modifications. Indeed, we observed that this band was immunoreactive against anti-SUMO2/3 antibodies (Figure 5A), indicating that a significant fraction of ERα-associated TOP2B was SUMOylated. Based on this, we turned our attention towards ZATT (zinc finger protein 451 associated with TDP2 and TOP2), the only TOP2B SUMO-ligase known so far.^47^ We found that ZATT also interacted with ERα, as proven by reciprocal endogenous co-immunoprecipitation experiments (Figure 5A-B), and that its depletion reduced the SUMOylation levels of ERα-bound TOP2B (Figure 5C). Together, these results indicate that ZATT associates with the ERα-TOP2A-TOP2B hub, within which it SUMOylates TOP2B.

**Figure 5.**
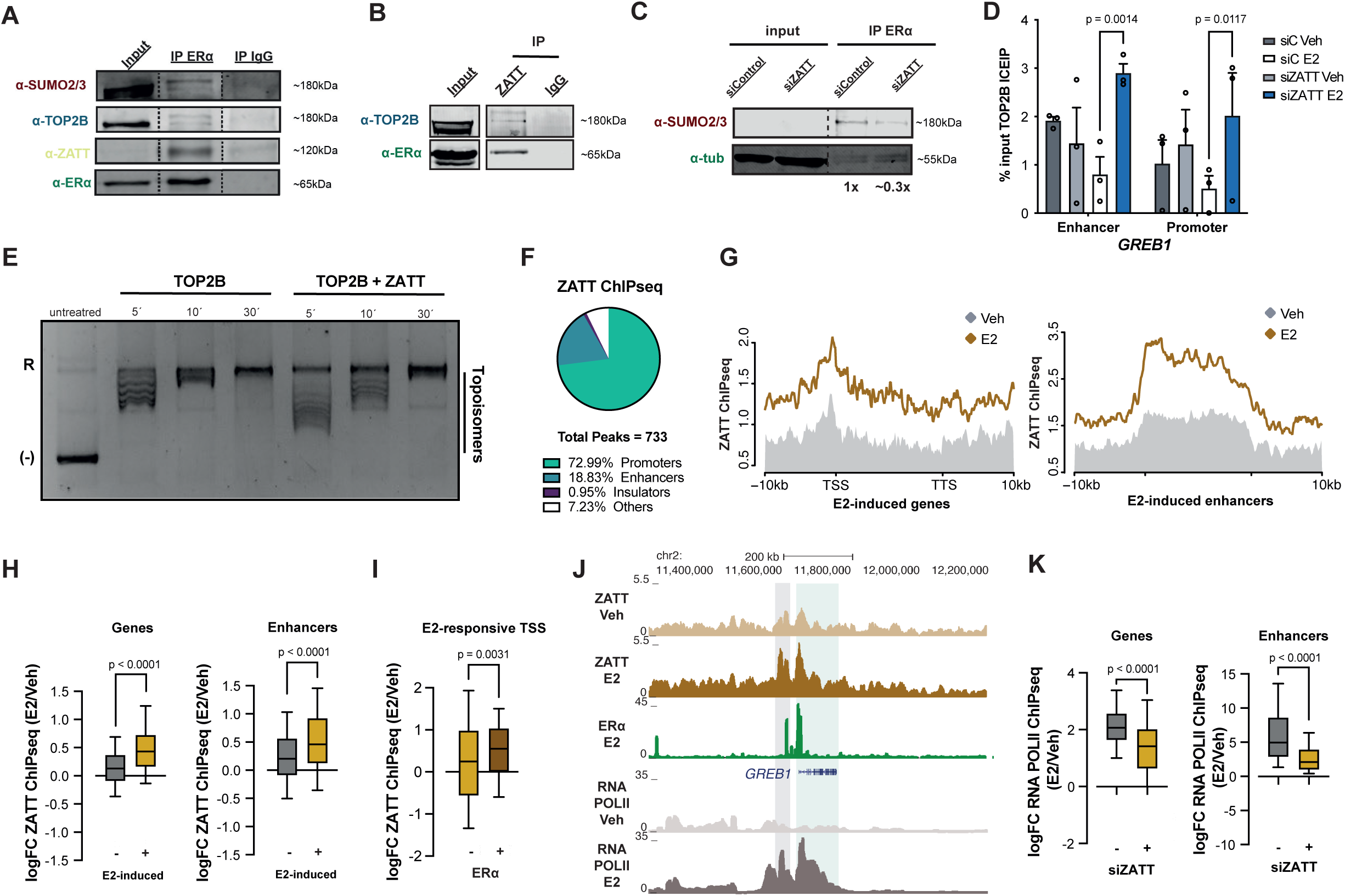
ZATT inhibits TOP2B activity to facilitate the estrogen response. (A) Immunoblot of SUMO2/3 (first panel), TOP2B (second panel), ZATT (third panel) and ERα (forth panel) in extracts of MCF7 cells (Input) and immunoprecipitated material (IP) using anti-ERα and control IgG antibodies, as indicated. (B) Immunoblot of TOP2B (top) and ERα (bottom) in extracts of MCF7 cells (Input) and immunoprecipitated material (IP) using anti-ERα and control IgG antibodies, as indicated. (C) Immunoblot of SUMO2/3 (top) and tubulin control (bottom) of extracts (Input) and anti-ERα immunoprecipitated material (IP) from MCF7 cells transfected with non-targeting control (siControl) or ZATT-specific (siZATT) siRNAs. Relative quantification of SUMO2/3 signal related to tubulin control is indicated below. (D) ICEIP qPCR experiments showing TOP2B activity at *GREB1* enhancer and promoter in hormone-depleted MCF7 cells transfected with non-targeting control (siC) or ZATT specific (siZATT) siRNAs in untreated control conditions (vehicle, Veh) or upon E2 treatment (10 nM, 45 min). Mean + SEM of at least three independent experiments is shown. Statistical analysis was performed using Two-way ANOVA. (E) Gel electrophoresis analysis of negatively supercoiled pBR322 plasmid incubated with recombinant TOP2B or TOP2B and ZATT for the indicated time. The positions of relaxed (R), negatively supercoiled (-) and topoisomers of pBR322 plasmid are indicated. (F) Annotation of ZATT ChIPseq peaks at promoters, enhancers and insulators. (G) Metaplot representation of average ZATT ChIPseq enrichment (RPGC) along E2-responsive genes (left) and enhancers (right) in hormone-depleted MCF7 cells transfected, in untreated control conditions (vehicle, Veh) or upon E2 treatment (10 nM, 45 min). (H) Average fold change (median of log2) of ZATT ChIPseq enrichment (RPGC) upon E2 treatment (45 min, 10 nM) at unresponsive and E2-induced genes (left) and enhancers (right). Boxplots represents 10-90 percentile and median of the signal. Statistical analysis was performed using Wilcoxon test. (I) As in “H” at E2-induced promoters that recruit ERα (logFC > 1.5) or not, as indicated. (J) Genome browser view of ZATT, ERα and RNA Pol II ChIPseq normalized signals (RPGC) at the *GREB1* regulatory landscape in the conditions of “H” and “I”. Grey and green boxes indicate *GREB1* enhancer (as determined by previous ERα ChIAPET experiments ^6^) and gene, respectively, (K) Average fold change (median of log2) of RNA Pol II CHIPseq signal (RPGC) upon E2 treatment (45 min, 10 nM) at E2-induced genes (left) and enhancers (right) in hormone-depleted MCF7 cells transfected with non-targeting control (siC) or ZATT specific (siZATT) siRNAs. Boxplots represents 10-90 percentile and median of the signal. Statistical analysis was performed using Wilcoxon test.

We then tested whether ZATT could be responsible for the downregulation of TOP2B activity that takes place in response to estrogen, and started by performing TOP2B ICE-IP qPCR analysis at the *GREB1* locus. Indeed, ZATT depletion abolished the downregulation of TOP2B activity observed upon E2 stimulation (Figure 5D), indicating that ZATT is necessary for the negative regulation of TOP2B activity in the context of ERα signaling. Since ZATT interacts with, and SUMOylates, TOP2B, we then wondered whether it could directly inhibit TOP2B activity, and whether this could depend on TOP2B SUMOylation, for which we performed *in vitro* plasmid relaxation assays. As can be observed in Figure 5E, the presence of recombinant ZATT protein significantly reduced, on its own, the capacity of TOP2B to relax (-) supercoiled pBR322 plasmid. In addition, when the experiment was performed under SUMOylation-competent conditions, we found no further inhibition (Figure S5A), even though a significant fraction of TOP2B was indeed SUMOylated (Figure S5B). Collectively, these results indicate that ZATT has the intrinsic capacity to inhibit TOP2B, suggesting that it may be a direct regulator of its activity during the estrogen response.

To test this possibility, we next characterized ZATT binding dynamics and its impact on the estrogen-dependent transcriptional response. First, we performed ZATT ChIP-seq experiments and studied its genome-wide distribution before and after E2 treatment. In general, ZATT was mostly located at promoter regions, yet exhibiting certain enrichment at enhancer regions (Figure 5F). Interestingly, we observed that, upon E2 stimulation, ZATT was recruited to estrogen-induced genes and enhancers (Figure 5G), and that this was specific when compared to non-responsive elements (Figure 5H), which is in line with an inhibitory function on TOP2B activity. Indeed, decreased TOP2B ICE-seq signal strongly correlated with ZATT recruitment at ERα-mediated enhancer contacts, even more so than with ERα or TOP2A (Figure S5C). Interestingly, when classifying estrogen-induced protein-coding genes depending on whether they displayed an ERα-binding site at their promoter or not, we observed that ZATT was differentially recruited to those bound by ERα (Figure 5I), and indeed, the level of ZATT and ERα recruitment strongly correlated (Figure S5C). Again, this can be clearly observed at the *GREB1* locus as an illustrative example (Figure 5J), where ZATT ChIP-qPCR analysis further validated the observed recruitment, as well as the specificity of the antibody (Figure S5D and S5E). Overall, these results agree with that the downregulation of TOP2B being directly mediated by ZATT (Figure 5D), while further supporting the idea that it particularly takes place at ERα-bound sites (Figure 4A).

Finally, to address whether ZATT-mediated downregulation of TOP2B activity facilitates the estrogen response, we performed RNA Pol II ChIP-seq experiments in ZATT-depleted cells. Interestingly, we observed a significant and specific reduction of E2-induced RNA Pol II recruitment to estrogen-induced genes and enhancers upon ZATT depletion (Figure 5K; see also an illustrative example in Figure S5F). Altogether, the results presented uncover the local ZATT-mediated downregulation of TOP2B activity as a molecular mechanism to facilitate the estrogen-induced transcriptional response through the stimulation of contacts between distal chromatin regulatory regions.

## DISCUSSION

DNA supercoiling has been traditionally considered an undesired byproduct of transcription that is resolved by DNA topoisomerases to generally favor gene expression. However, this view is changing in recent years, and topoisomerases are being assigned relevant regulatory functions.^23,30,34^ In these cases, topoisomerases are normally believed to act as activators of transcription, while their mechanism of action still remains poorly characterized. Studying TOP2 functions during the estrogen-induced transcriptional response, we have uncovered a repressive role of TOP2B that operates by removing DNA supercoiling and restraining regulatory chromatin contacts.

In our model (Figure 6), TOP2B activity at estrogen-responsive elements would remove the topological stress generated under basal transcription conditions. Upon exposure to estrogen, ERα, TOP2A and ZATT are recruited to these regions, leading to a downregulation of TOP2B activity that results in an accumulation of (-) DNA supercoiling that would favor regulatory chromatin interactions and stimulates the estrogen response. In this scenario, the regulatory regions would have an intrinsic tendency to interact, which would be normally kept in check by continuous TOP2B-mediated removal of (-) supercoiling. The feedback loop of more (-) supercoiling leading to more transcription that generates more (-) supercoiling and so on, would allow a rapid response upon TOP2B inhibition. It is worth noting that (-) supercoiled DNA can provide a particular topological context in which distal chromatin contacts are favored while maintaining permissive conditions for transcription, i.e., by facilitating promoter melting, the advance of RNA Pol II and/or nucleosome remodeling.^48–52^ Importantly, this model provides an explanation for the regulatory activity of TOP2B based on its well-established action on transcription-associated topological stress without invoking additional functions, such as, for example, the generation of DNA breaks, which have been proposed to induce estrogen-dependent gene transcription.^33^ This apparent discrepancy may be explained by the previous difficulty to measure localized paralog-specific TOP2 activity, which we have overcome with the development of ICE-seq and ICE-IP qPCR methodologies, and if DNA breaks are considered an accidental consequence, rather than the cause, of transcriptional upregulation.

**Figure 6.**
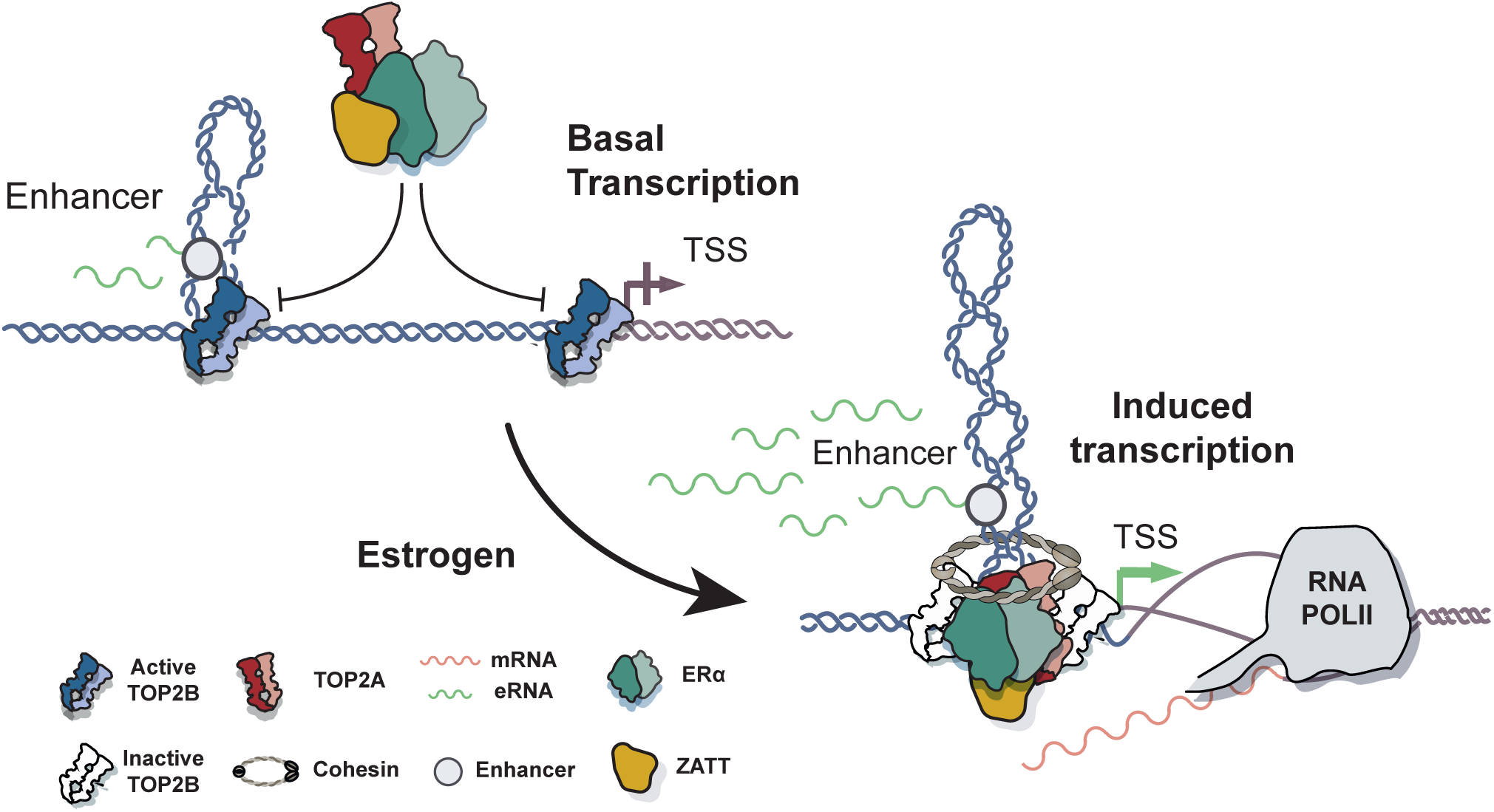
Model of topological control of the estrogen response by the regulation of TOP2B activity. Under basal transcription conditions TOP2B removes the generated (-) supercoiling. In response to estrogen, ERα, TOP2A and ZATT recruitment to E2-responsive elements triggers a downregulation of TOP2B activity that results in an accumulation of (-) supercoiling that facilitates ERα-mediated chromatin contacts and induces transcription.

Our results argue that physiological downregulation of TOP2B activity at estrogen-responsive elements depends on ERα, TOP2A and ZATT. It is difficult to unambiguously assign a direct function in TOP2B inactivation to ERα, since, being the molecular trigger of the entire pathway, all downstream events are expected to be affected. We, however, believe that the correlation of its recruitment with the reduction in TOP2B activity, together with the co-occupancy and physical interactions with TOP2A, TOP2B and ZATT, strongly suggest that ERα is indeed directly involved in TOP2B downregulation and the topological control of the estrogen response. Regarding TOP2A, our findings suggest a dual role in stimulating the estrogen response, both, by facilitating estrogen-induced transcription through the removal of associated topological stress, as previously suggested,^29,30,32^ and by participating in TOP2B inhibition through a non-catalytic function that is particularly apparent at enhancer regions, where TOP2A is recruited but its activity is only mildly increased. Finally, our results suggest that ZATT is likely the effector that directly inhibits TOP2B activity. We show that ZATT has the *in vitro* ability to inhibit TOP2B relaxation activity in a SUMOylation-independent manner. We cannot however rule out the possibility that, in a cellular context, ZATT-mediated SUMOylation, either of TOP2B or some accessory factor, may enhance or mediate TOP2B inhibition. The specific interplay between TOP2 paralogs and ZATT in transcriptional regulation is an interesting matter for future investigation.

Our model links DNA supercoiling with 3D genome architecture, a phenomenon long-studied in bacteria^53,54^ that is currently emerging in eukaryotes.^20,55,56^ How can transcription-derived DNA supercoiling facilitate regulatory chromatin contacts? On the one hand, DNA supercoiling, in the form of superhelical structures, may simply favor a 3D conformation of the chromatin fiber that physically reduces the distance between regulatory elements, as previously shown in vitro.^18^ On the other hand, a non-mutually exclusive possibility, would be that DNA supercoiling “pushes’’ the cohesin ring along the chromatin fiber, thereby facilitating DNA loop extrusion.^20,21^ This is in line with recent reports highlighting the relevant interplay between transcription and cohesin dynamics to shape genome organization,^16,57–59^ and enhancer function in particular.^14,15,60,61^ In this sense, we observed that cohesin is accumulated at ERα-mediated contact points upon TOP2B depletion, suggesting that these supercoiling-mediated contacts would be also associated with the cohesin complex, either as *de novo* loading to stabilize the interactions, or as part of a supercoiling-driven loop extrusion mechanism. ERα binding, either on its own or in conjunction with RNA Pol ll,^16,17^ could also help to shape regulatory contacts by representing an impediment for the spreading of supercoiling and/or cohesin movement, which would explain its preferential location at these contact points. The combination of increased supercoiling levels with the appearance or strengthening of barriers for its spreading would constitute an ideal scenario for the rewiring of long-range chromatin interactions.

The physiological modulation of DNA supercoiling levels would represent a novel regulatory layer that may not be limited to estrogen signaling, but could be a common feature of transcriptional circuits involving a significant rewiring of chromatin contacts. This is in line with the reported role of ZATT in androgen receptor-dependent gene activation,^62^ which could, in the light of our findings, also operate through TOP2B regulation. Furthermore, even other cellular processes of genome dynamics with a strong topological and architectural component, such as for example replication, could also rely to some degree on an integrated control of DNA supercoiling and chromatin organization, highlighting the potential relevance of DNA topoisomerases as master regulators of genome dynamics. Given that combined treatments with hormonal and TOP2 inhibitors are common practice in ER-positive breast and ovarian cancers, the unravelling of the topological regulation of estrogen signaling may be of therapeutic importance.

## ACKNOWLEDGMENTS

We thank the Genomics and Microscopy Core facilities at CABIMER and the Genomics and Confocal Microscopy Units ant CNIO. Computational analyses were run on the High-Performance Computing cluster provided by the Centro Informático Científico de Andalucía (CICA). The F.C.-L. laboratory is supported by grants from MCIN/AEI/10.13039/501100011033 and ERDF “A way of making Europe” (SAF2017-89619-R and PID2020-119570RB-I00) and the European Research Council (ERC Consolidator Grant, ERC-CoG-2014-647359 TOPOmics). The G.M.-Z. laboratory is supported by grants from MCIN/AEI/10.13039/501100011033 (PID2021-127432N) and (CNS2022-135600). A.A is supported by grants from MCIN/AEI/10.13039/501100011033 and ERDF “A way of making Europe” (PID2019-104270GB-I00/BMC) and the European Research Council (ERC Advanced Grant, ERC-AdG-2014-669898 TARLOOP). Work in the R.S.W laboratory is supported by the US National Institute of Health Intramural Program, US National Institute of Environmental Health Sciences (NIEHS) 1Z01ES102765. J.T.-B. was supported by a fellowship from Ministerio de Educación, Cultura y Deporte (FPU15/03656). M.M.-S was supported by a fellowship from Agencia Estatal de Investigación (SEV-2015-0510-19-1, PRE2019-090967). G.M.-Z. was supported by Asociación Española Contra el Cáncer (POSTD18021MILL) and a fellowship from “la Caixa” Foundation (ID 100010434) and from the European Union’s Horizon 2020 research and innovation programme under Marie Skłodowska-Curie grant agreement No. 847648. The fellowship code is LCF/BQ/PR21/11840007.

## AUTHOR CONTRIBUTIONS

Conceptualization: J.T.-B., G.M.-Z. and F.C.-L. Methodology: J.T.-B., G.M.-Z. and F.C.-L. Formal analysis: J.T.-B. Investigation: J.T.-B., M.M.-S., L.L.-H. and G.M.-Z. Resources: A.J.V., M.G.-D. and R.S.W. Writing - Original Draft: G.M.-Z and F.C.-L. Writing - Review & Editing: J.T.-B., A.A., G.M.-Z and F.C.-L. Supervision: A.A., G.M.-Z and F.C.-L. Funding acquisition: A.A., G.M.-Z. and F.C.-L.

## DECLARATION OF INTERESTS

Authors declare no competing interest.

## SUPPLEMENTARY FIGURE LEGENDS

**Supplementary Figure 1.**
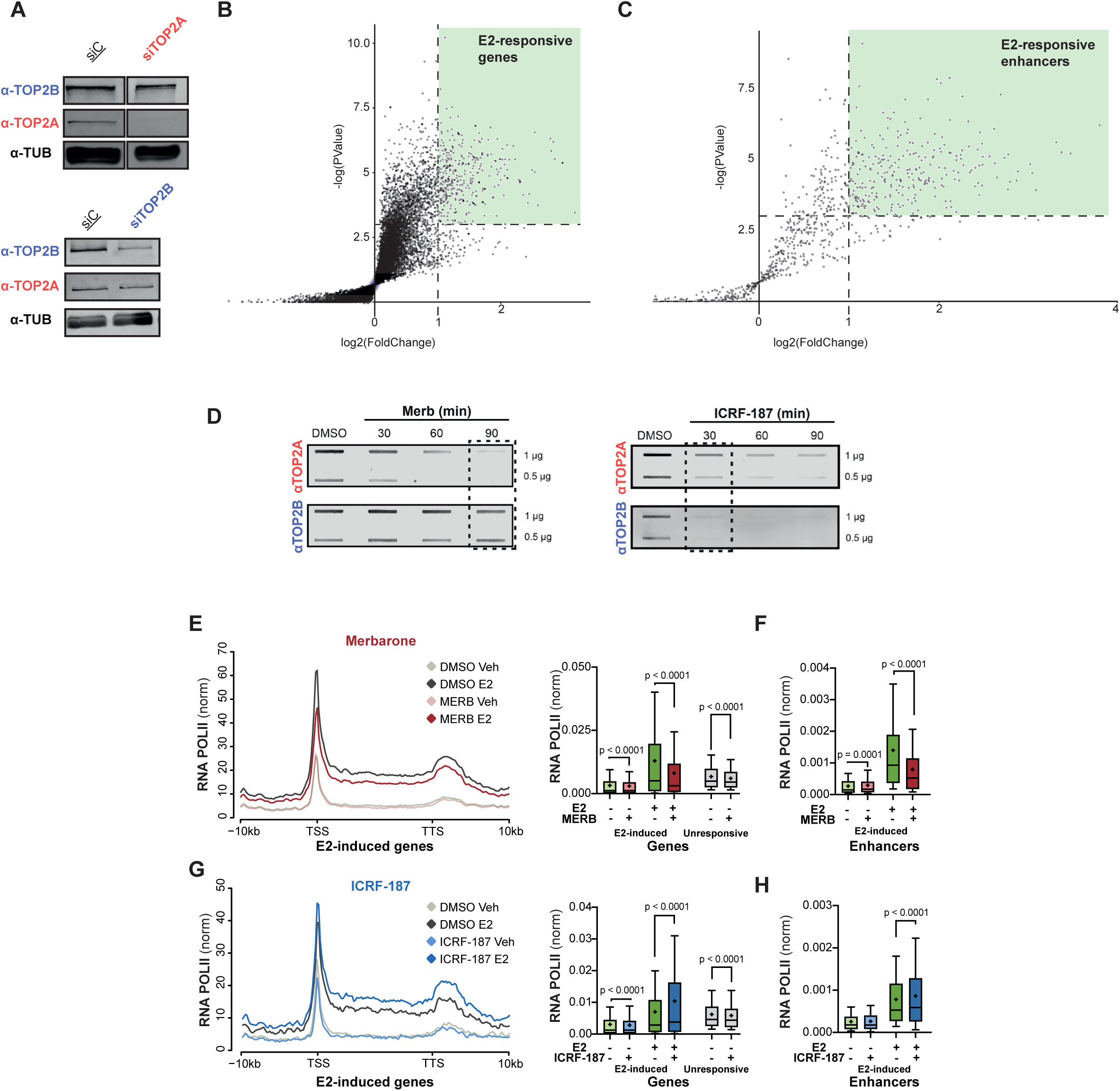
(A) Representative image TOP2A and TOP2B protein levels after siRNA depletion (72h) as determined by Western blottting. (B) Representation of RNA POLII enrichment at annotated protein-coding genes and enhancers upon E2 treatment (10 nM, 45 min). Counts are normalized by total number of reads and spike-in factor. Fold-change (log2) of 4 independent experiments and p-value (-log), as determined by t-test, is shown. The green square indicates log2FC > 1; p-value < 0.05, as used for selection of E2-induced genes. (C) As in “B” for E2-induced enhancers, identified in previous GRO-seq experiments. (D) Representative image of ICE experiments showing TOP2Accs (top) and TOP2Bccs (bottom) in hormone-depleted MCF7 cells treated with etoposide (100 µM for 30 minutes) and their reduction by pre-treatment with merbarone (100 µM) or ICRF-187 (100 µM) for the indicated times. Two concentrations of genomic DNA are shown to illustrate the quantitative range of the signal. Conditions showing preferential inhibition are highlighted (squares). (E) Metaplot (left) and quantification (right) of average RNA Pol II ChIPseq enrichment, represented as normalized reads per genome coverage (RPGC), along E2-responsive genes in hormone-depleted MCF7 cells pre-treated with merbarone (100 µM, 90 min) in untreated control conditions (vehicle, Veh) or upon E2 treatment (10 nM, 45 min). Quantification of average RNA Pol II ChIPseq enrichment, represented as normalized reads per genome coverage (RPGC) at shown E2-responsive genes. (F) Quantification of average RNA Pol II ChIPseq enrichment, represented as normalized reads per genome coverage (RPGC) at E2-responsive enhancers. Other details as in “E”. (G and H) As in “E” and “F” in cells pre-treated with ICRF-187 (100 µM, 90 min).

**Supplementary Figure 2.**
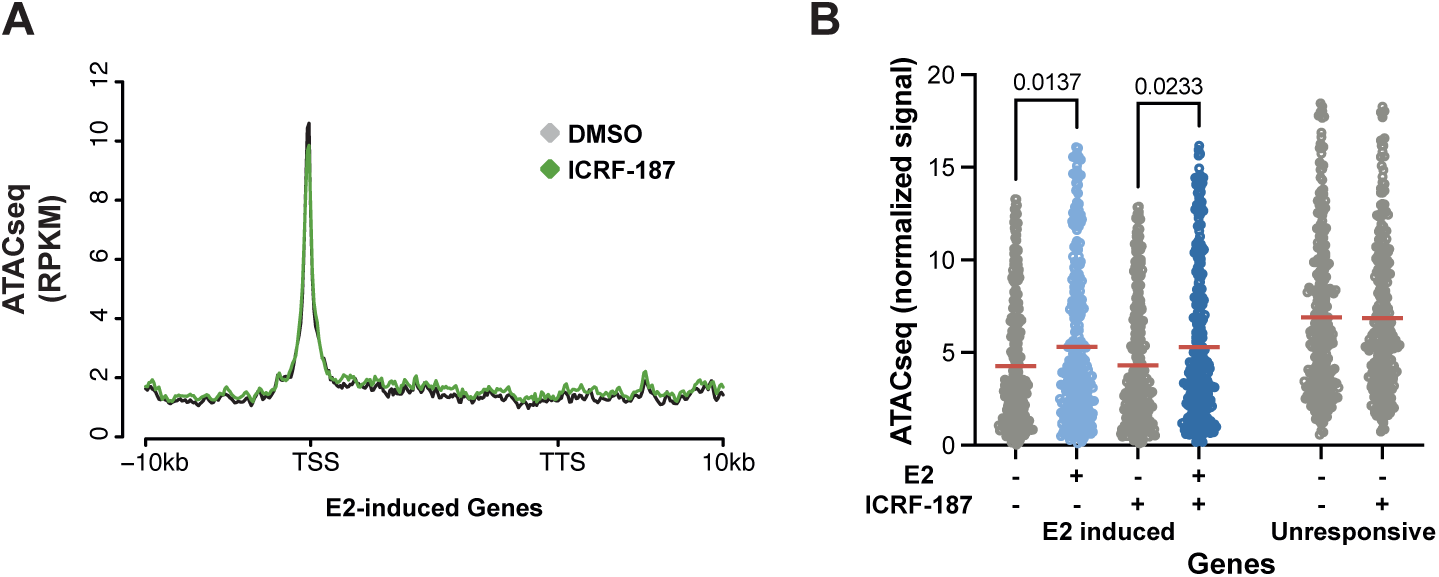
(A) Metaplot representation of ATACseq profiles (RPKM) at E2-responsive genes upon ICRF-187 treatment (100 µM, 30 min). Transcribed regions +/- 10 kb are shown. (B) Quantification of of ATACseq signal at E2-induced and unresponsive protein-coding genes and upon E2 and ICRF-187 treatment, as indicated. Mean is shown in red bars. Statistics by One-way ANOVA with Turkey post-test.

**Supplementary Figure 3.**
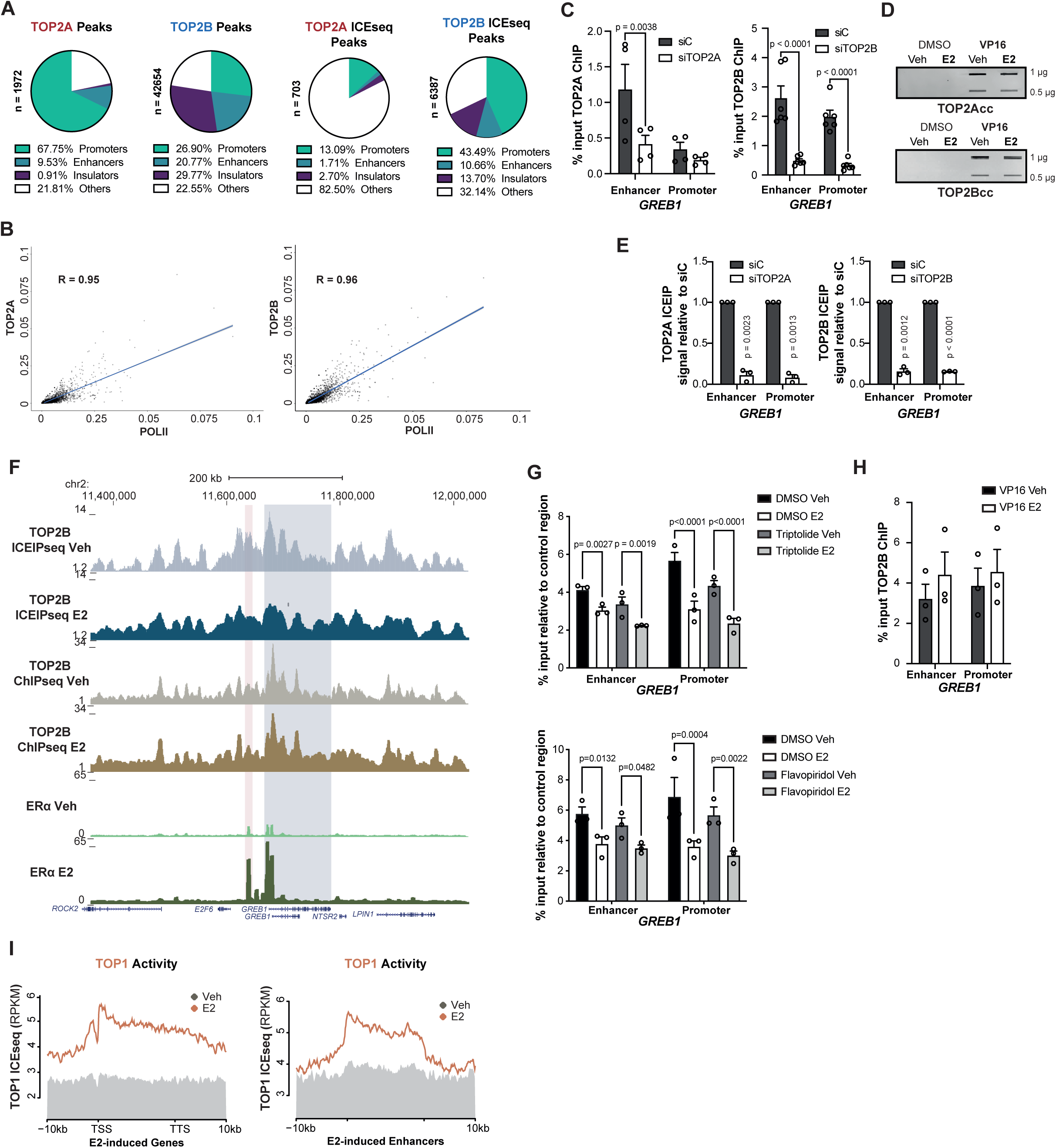
(A) Genome annotation of identified TOP2A and TOP2B ChIPseq and ICEseq peaks. (B) Spearman correlation between RNA POLII and TOP2A or TOP2B ChIPseq signal at transcribed regions. (C) TOP2A and TOP2B ChIP qPCR signal at *GREB1* enhancer and promoter upon TOP2A and TOP2B depletion by siRNA (72 h), as indicated. Statistics by Two-Way ANOVA. (D) Representative image of ICE experiments showing TOP2Accs (top) and TOP2Bccs (bottom) in hormone-depleted MCF7 cells treated with etoposide (100 µM, 30 min) and/or E2 (100 µM, 30 min), as indicated. Two concentrations of genomic DNA are shown to illustrate the quantitative range of the signal. (E) TOP2A and TOP2B ICE-IP qPCR signal at *GREB1* enhancer and promoter upon TOP2A and TOP2B depletion by siRNA (72 h). Statistics by one-sample t-test. (F) Genome view of the *GREB*1 locus showing TOP2B ChIPseq and ICEseq profiles in response to E2 treatment. Red box corresponds to the *GREB1* enhancer and gray box to the annotated gene. (G) TOP2B ICEIP qPCR signal upon E2 treatment and flavopiridol (0.5 µM, 2h) and/or triptolide (10 µM, 1h) pre-treatment, as indicated. Average +/- S.E.M. of three independent experiments is shown. Statistics by Two-Way ANOVA. (H) TOP2B ChIP qPCR signal upon control and E2 treatment conditions VP16 treatment (100 µM, 30 min). Mean +S.E.M. of three independent experiments and significance by Two-Way ANOVA is shown. (I) Metaplot representation of TOP1 ICEseq profile (RPKM) at E2-responsive genes and enhancers upon E2 treatment (10 nM, 45 min). Transcribed regions +/- 10 kb are shown.

**Supplementary Figure 4.**
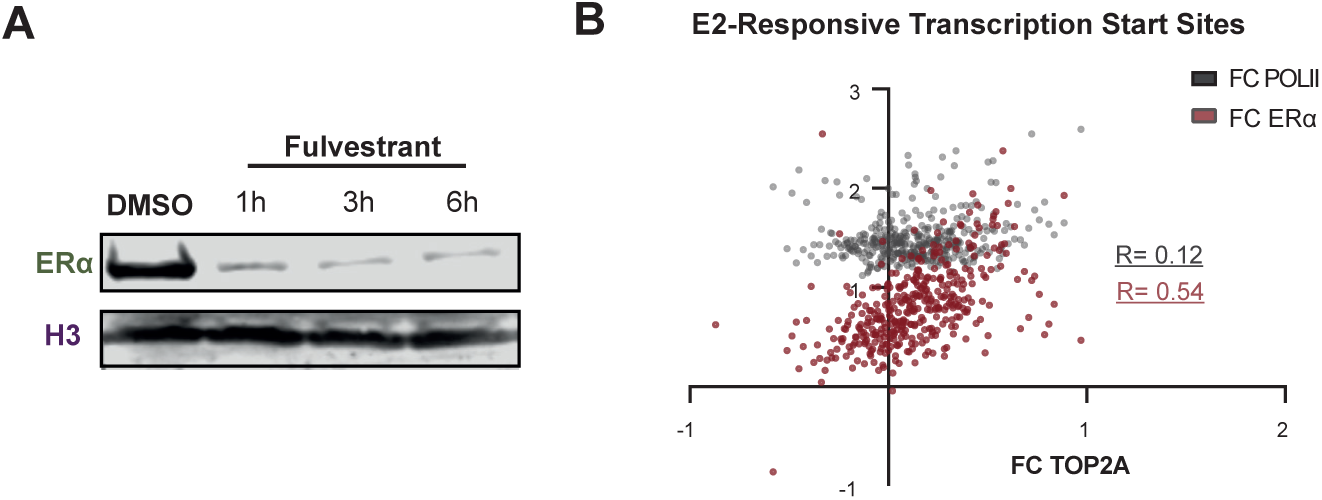
(A) Immunoblotting of ERα in control conditions (DMSO) and after the indicated times of 100 nM fulvestrant treatment. Histone H3 immunoblot is shown as loading control. (B) Spearman correlation between the recruitment (fold-change; FC) of TOP2A and ERα or RNA POLII upon E2 treatment at E2-responsive TSS (+/- 3 kb).

**Supplementary Figure 5.**
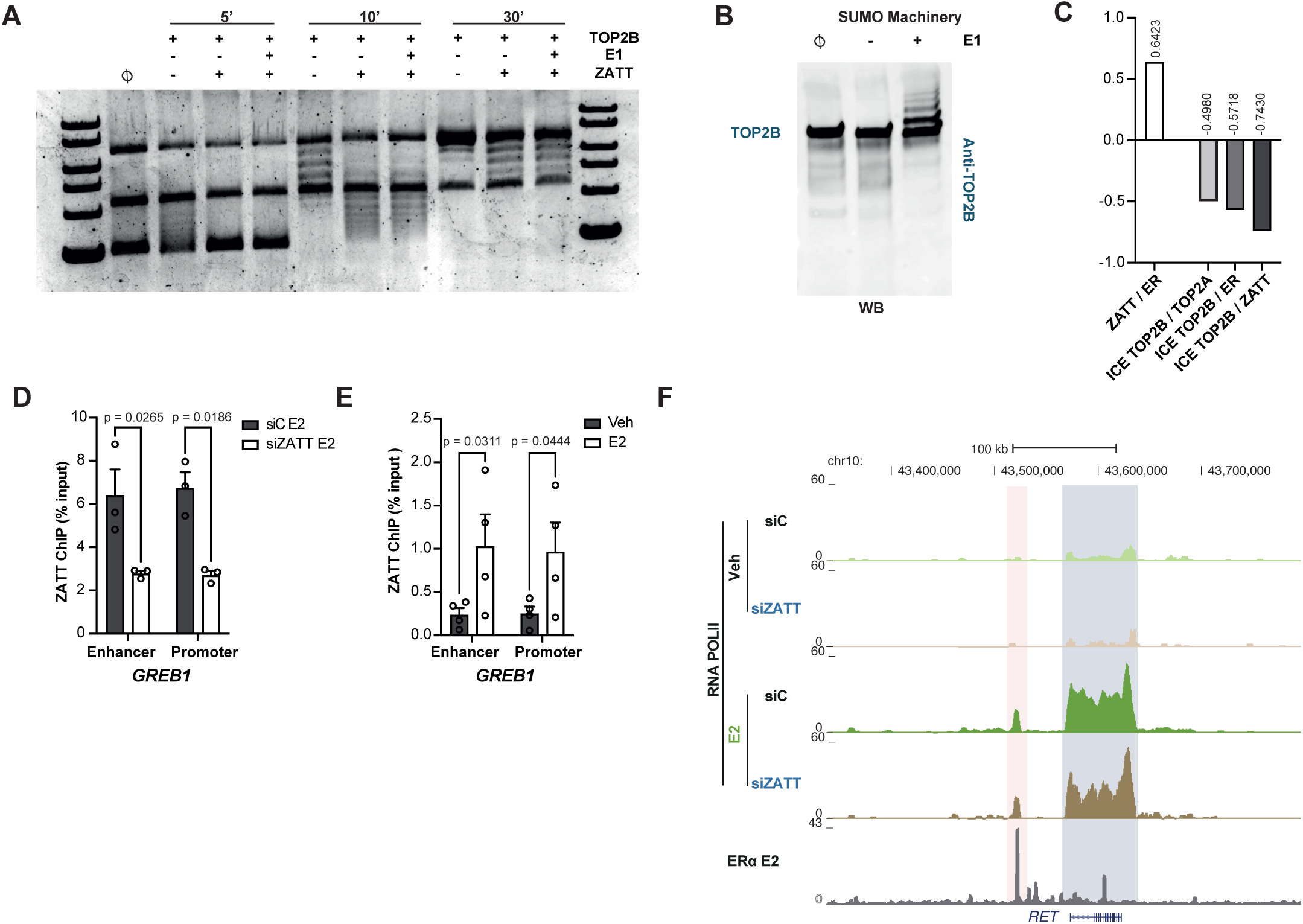
(A) Gel electrophoresis analysis of negatively supercoiled pBR322 plasmid incubated with SUMOylation components including or not, as indicated, the E1 ligase, TOP2B and/or ZATT for the indicated time. The positions of relaxed (R), negatively supercoiled (-) and topoisomers of pBR322 plasmid are indicated. (B) Representative image of an immunoblot detecting TOP2B in reactions containing SUMOylation components, TOP2B and ZATT, and, including or not, the E1 ligase, as indicated. (C) Spearman correlations between the indicated signals at enhancer contact points. (D) ZATT ChIP qPCR signal at the *GREB1* enhancer and promoter regions in hormone-depleted MCF7 cells transfected with non-targeting control (siC) or ZATT-specific (siZATT) siRNAs treated with E2 (10 nM, 45 min). (E) ZATT ChIP qPCR signal at the *GREB1* enhancer and promoter regions in hormone-depleted MCF7 cells in control conditions and upon E2 treatment (10 nM, 45 min). (F) Genome view of the *RET* locus showing ERα and RNA POLII ChIP-seq signal (normalized RPGC) in control conditions and in response to E2 treatment (10 nM, 45 min).

## STAR METHODS

### Key resources table

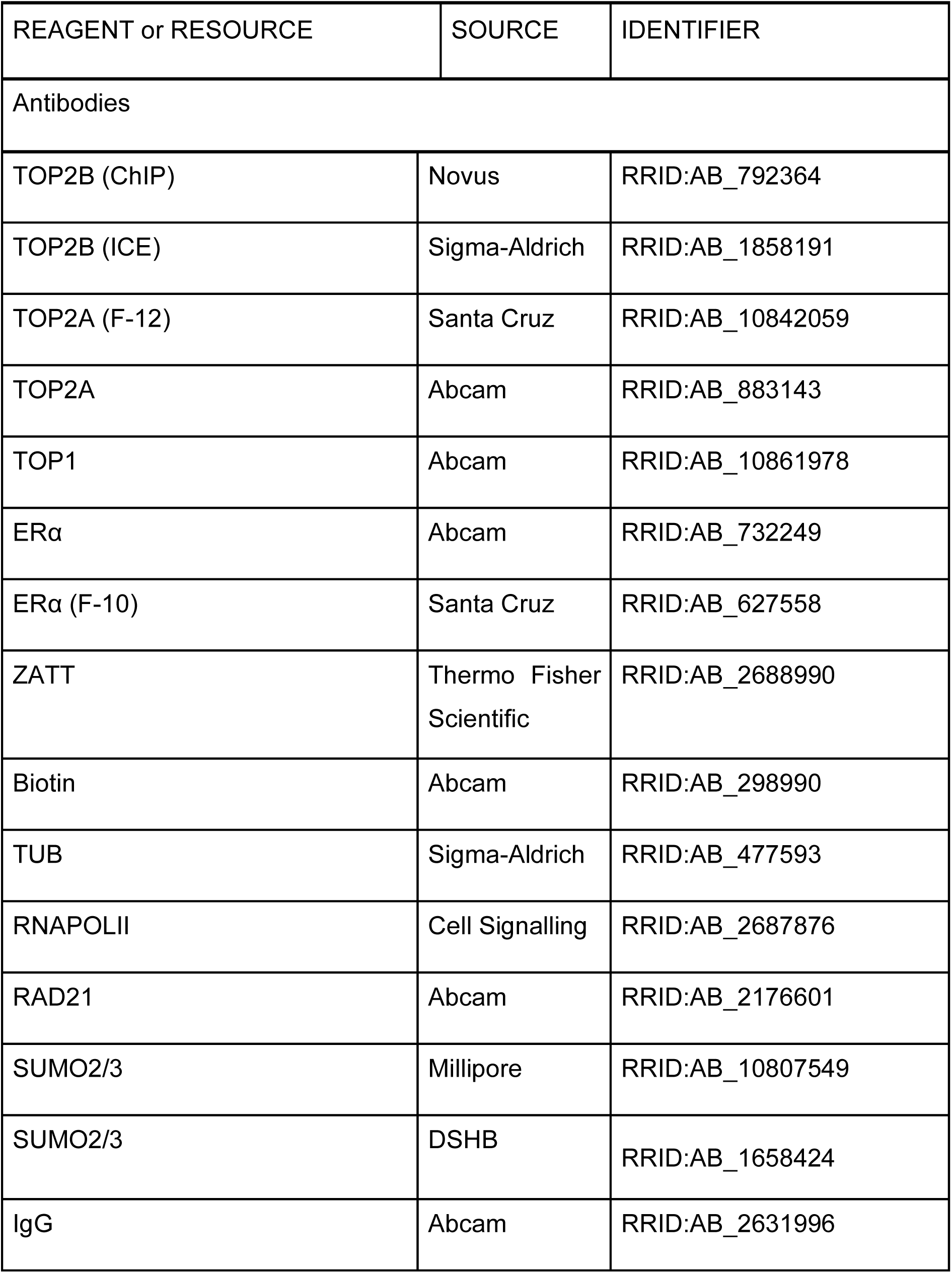
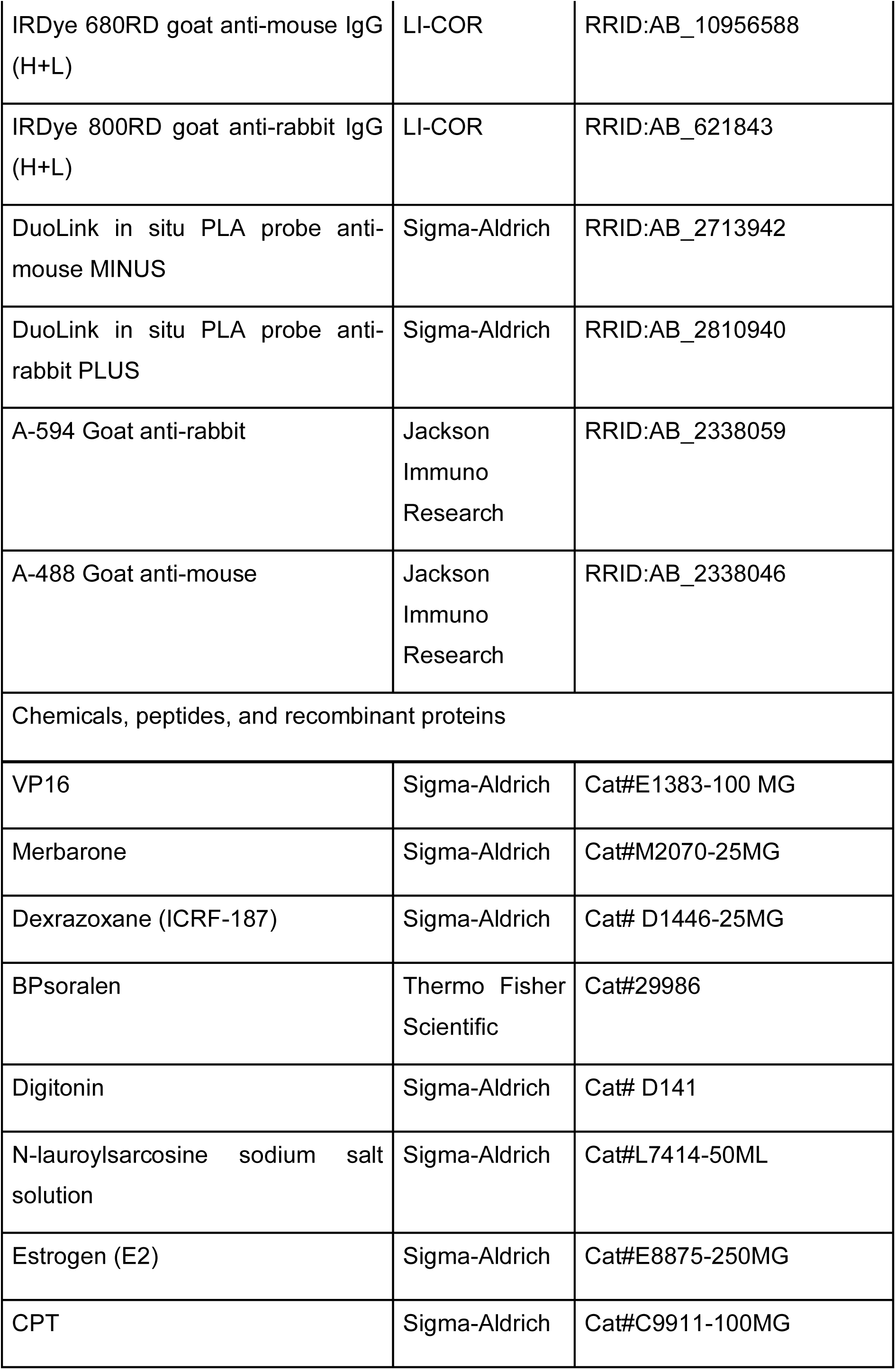
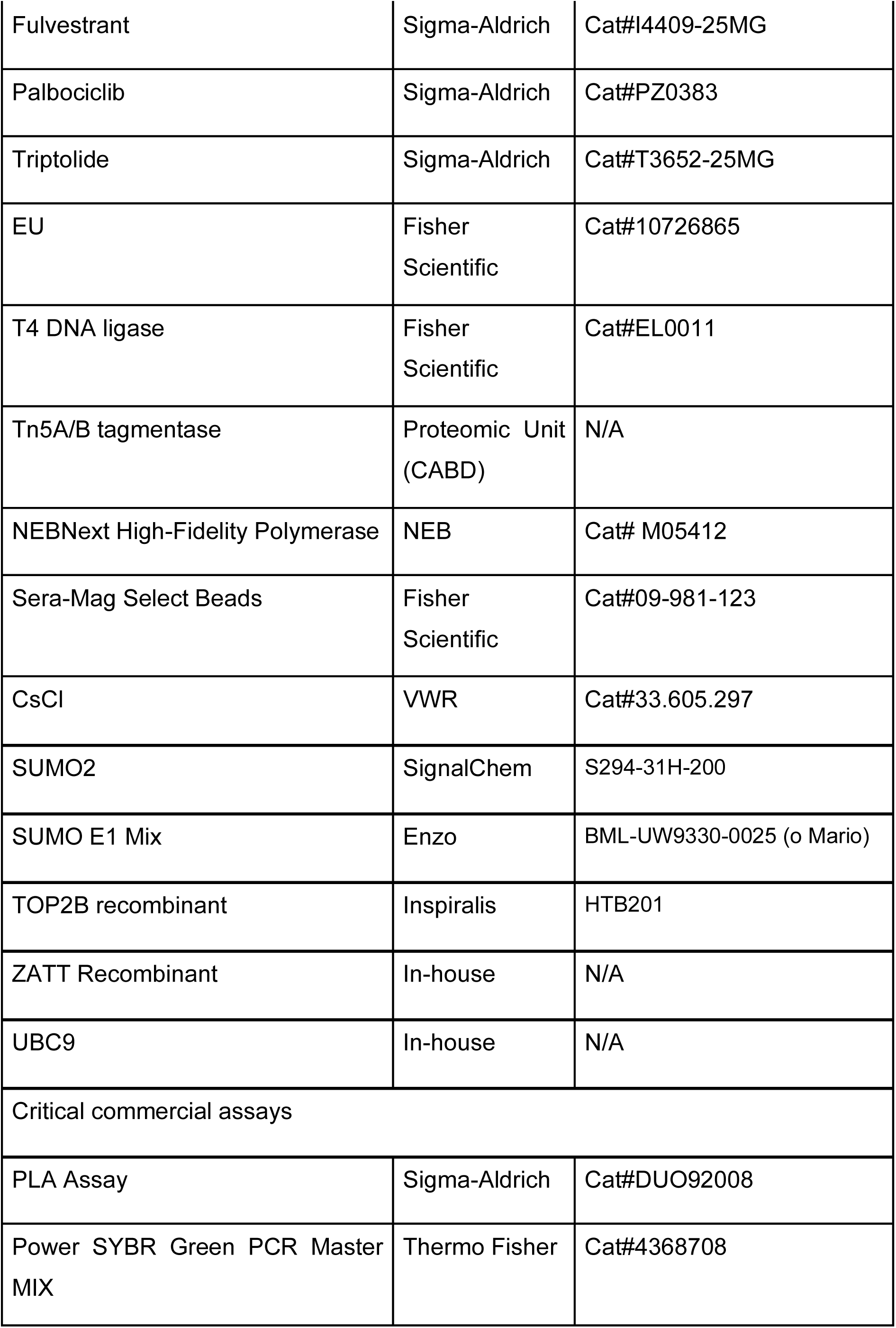
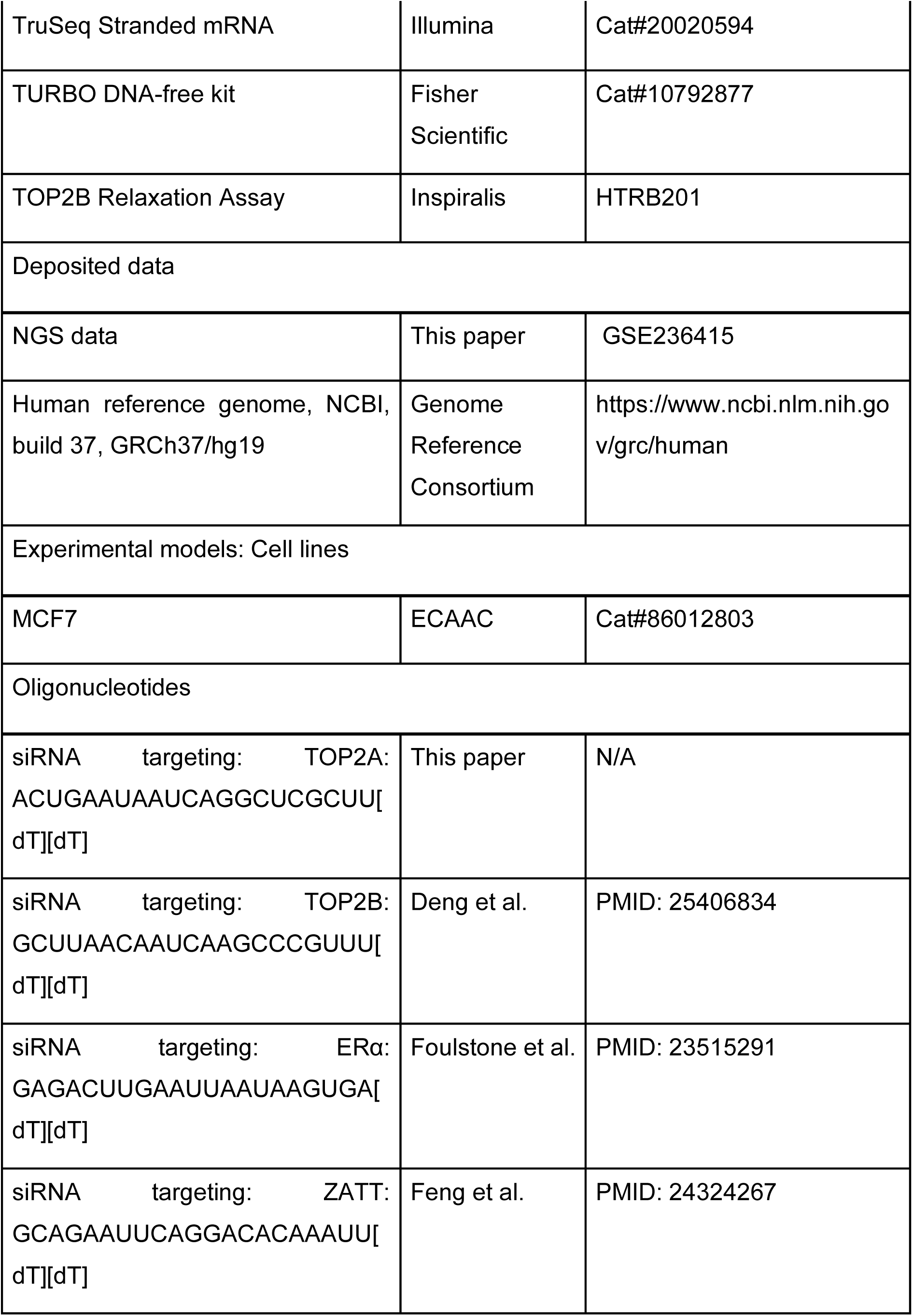
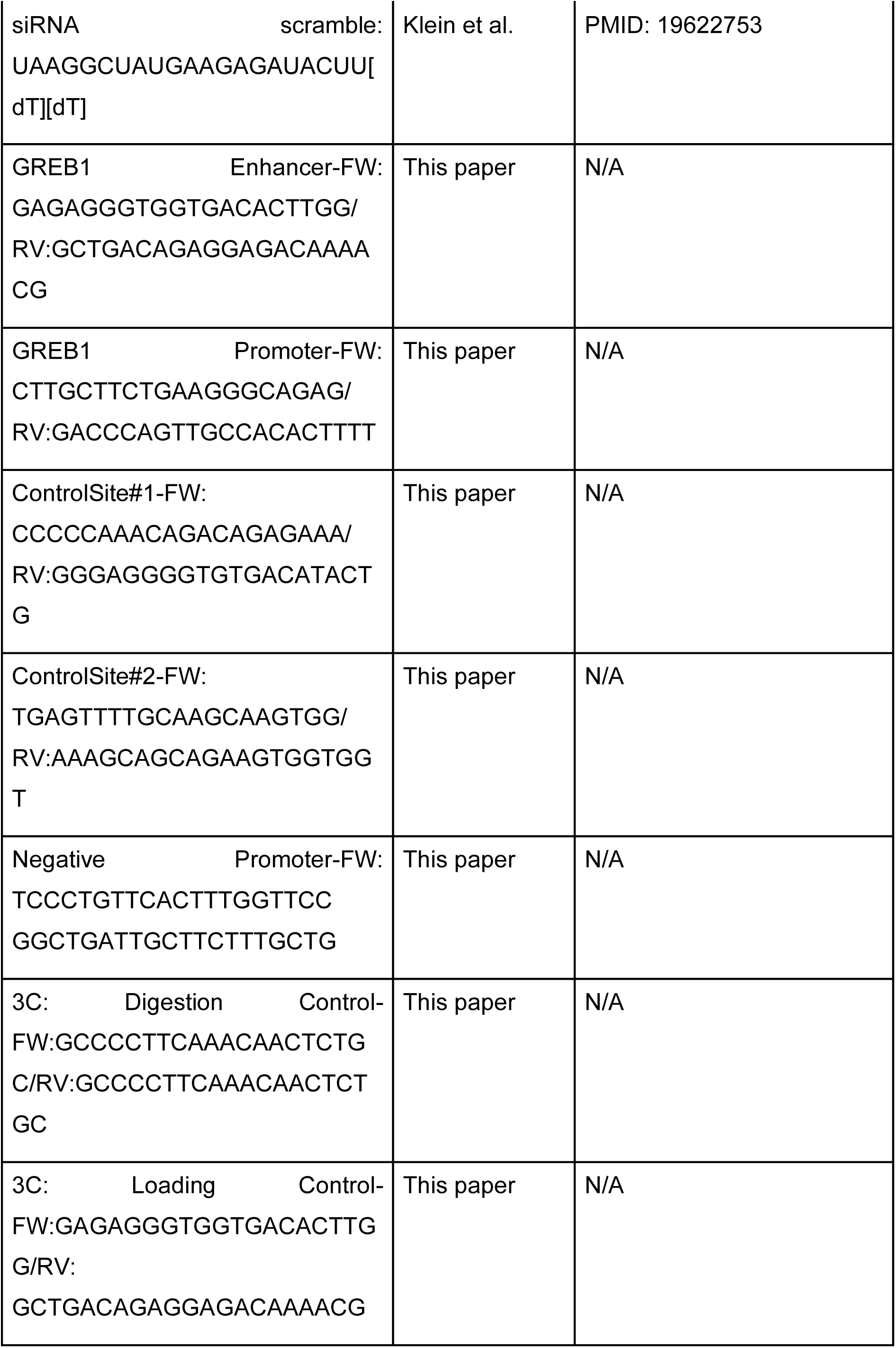
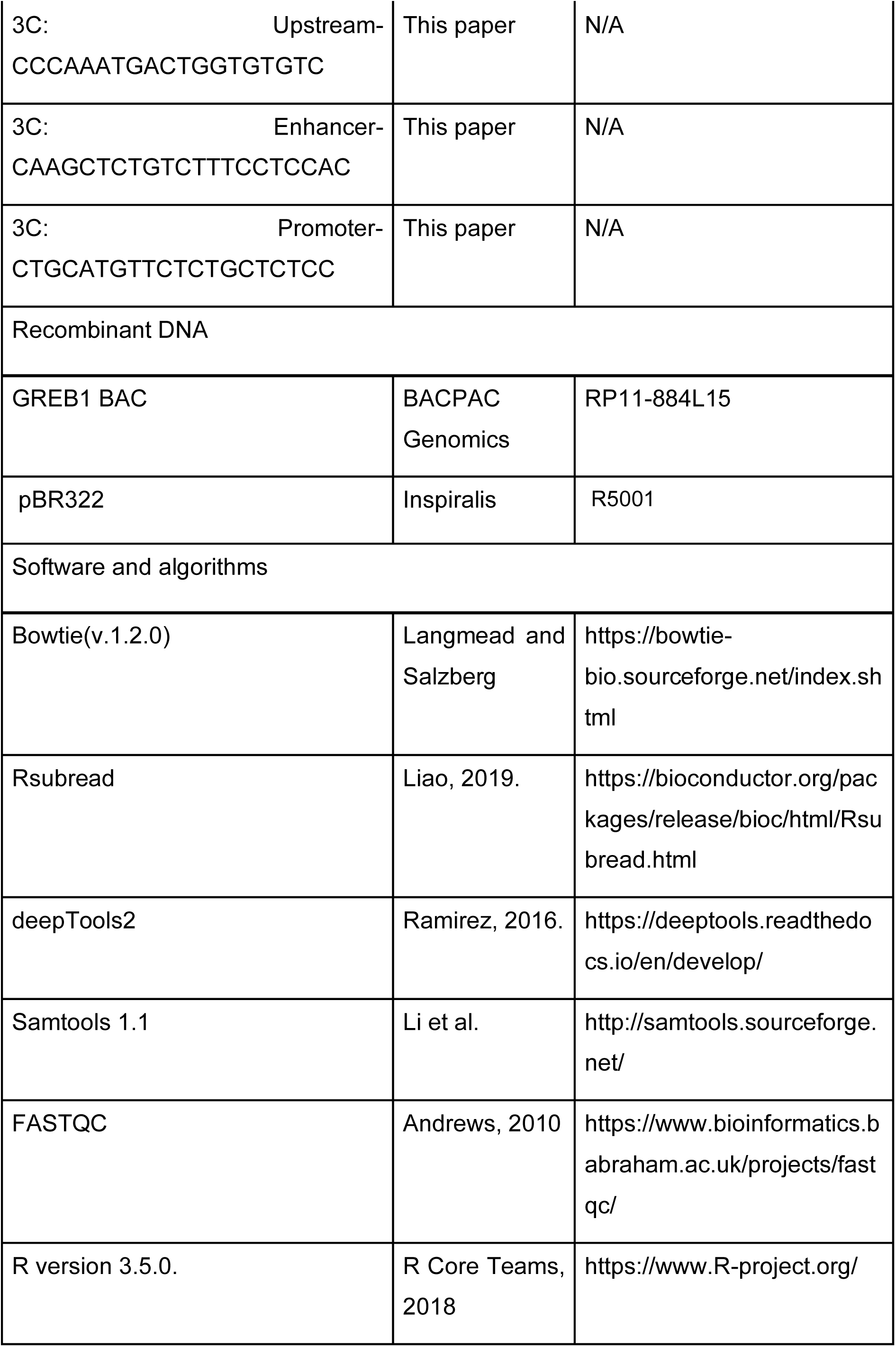
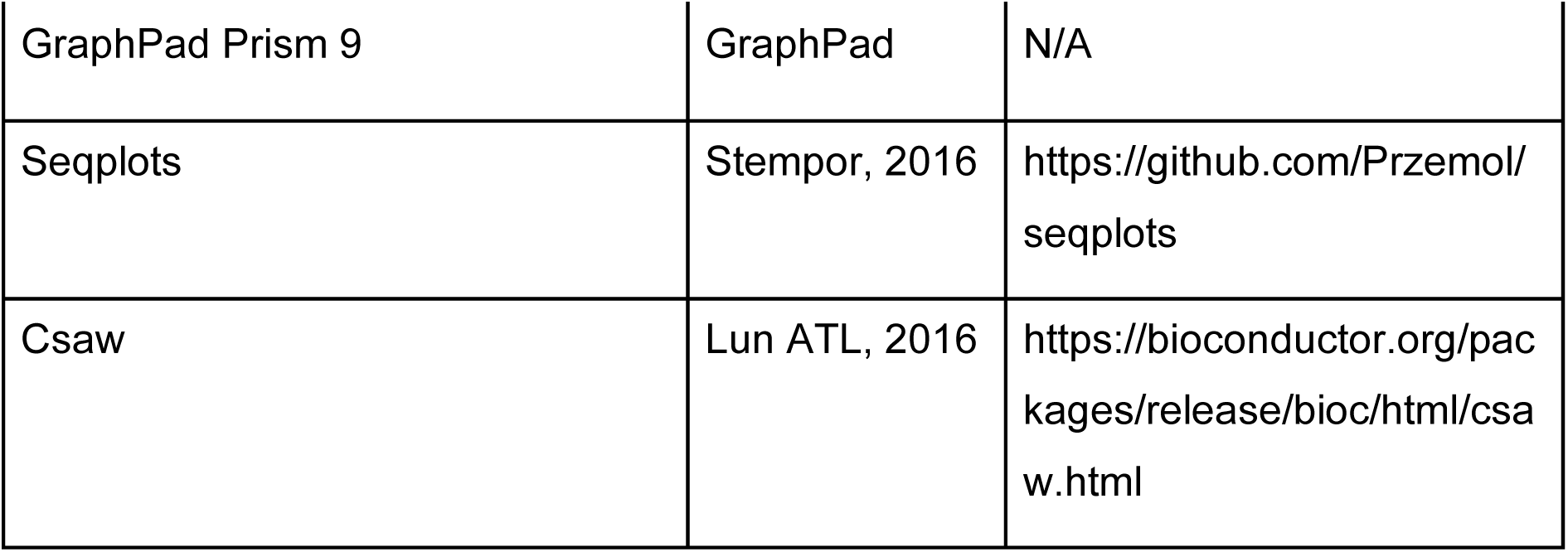

### Cell culture and treatments

MCF7 cells (ECACC, 86012803) were cultured in Minimum Essential Medium (MEM) with Earlés salts supplemented with 50 U/ml Penicillin, 50 µg/ml Streptomycin, 2 mM L-Glutamine, 10% FBS (Sigma), and non-essential amino acids. Cell maintenance was conducted in HEPA class-100 incubators (Thermo) at 37°C with 5% CO2. E. coli strains were grown in Luria-Bertani (LB) media with appropriate antibiotics at 37°C. Regular mycoplasma testing was performed using the MycoAlert PLUS Mycoplasma Detection Kit (Lonza).

For all experiments involving estrogen treatment, cells were deprived of hormones prior to estrogen induction (10 nM, 45 min). 16 hours after seeding cells in culture plates, the medium was replaced with MEM without phenol-red supplemented with 5% charcoal-stripped FBS (Sigma-Aldrich, F6765). Cells were incubated for 48 hours in these conditions before conducting the experiments. Control cells were treated with vehicle (EtOH). Transcription inhibition was achieved by pre-treating cells with 10 µM triptolide for 30 min. To inhibit topoisomerase activity, cells were preincubated with 100 µM Merbarone or 100 µM ICRF-187 for the indicated time.

Protein knockdowns were performed by transfecting cells with siRNAs 72h before conducting experiments. MCF7 were transfected using Lipofectamine RNAiMAX (Fisher Scientific, 12343563) following the standard protocol for a reverse transfection.

### *In situ* proximity ligation assay

The *in situ* PLA experiments were performed following the manufacturer’s protocol available from Millipore-Sigma. The following DuoLink *in situ* PLA probes and reagents were used: anti-mouse MINUS (DUO92004), anti-rabbit PLUS (DUO92002), and DuoLink Detection Reagents (DUO92008).

A total of 40,000 cells were seeded in 24-well plates and, after treatment, fixed in 4% PFA-PBS for 10 minutes at room temperature. Subsequently, cells were permeabilized with 0.2% Triton X-100 in PBS for 2 minutes. After three PBS washes, cells were subjected to 30-minute incubation at 37°C in a humidity chamber with a blocking solution, followed by a 90-minute incubation with primary antibodies at 37°C.

Coverslips were washed twice with buffer A and incubated with PLA probes for 1 hour at 37°C. After a single wash with buffer A, the ligation reaction was performed according to the standard protocol for 30 minutes at 37°C, followed by incubation with polymerase for rolling circle amplification during 90 minutes at 37°C. After two consecutive 10-minute washes with buffer B, cells were counterstained with DAPI (Sigma) and mounted with Vectashield mounting medium (VectorLabs). Images were captured using a Leica DM-6000B wide-field fluorescence microscope.

Foci located within cell nuclei were counted using Metamorph as follows: nuclei were selected based on DAPI staining and size (min width = 35 µm, max width = 150 µm and background level > 300 graylevels). Foci were then counted using “granularity” function with standard algorithms (min width = 0 µm, max width = 6 µm and background level > 200 gray levels). Figures display the average number of cells with more than 3 PLA foci from four independent experiments. Mean +/- SEM, with statistics by Chi-square with Yates correction, is shown.

### Chromatin Conformation Capture (3C)

3C experiments were conducted adapting the protocol described in ^63^. At least 5 x 10^6^ cells were fixed for 10 min at RT using 1% formaldehyde in pre-warmed non-supplemented MEM without phenol-red. Crosslinking was quenched by incubating with a final concentration of 150 mM glycine for 5 minutes at RT and 15 minutes at 4°C.

Next, cells were scraped into 2.5 ml of ice-cold PBS supplemented with the Complete protease inhibitor cocktail and centrifuged at 4°C and 800xg for 10 minutes. The resulting cell pellets were resuspended in 1 ml of cold hypotonic lysis buffer (10 mM Tris-HCl pH 8, 10 mM NaCl, 0.5% NP-40) supplemented with the Complete cocktail and incubated for 15 minutes on ice. After incubation, cells were homogenized and lysed on ice using a cell douncer, with a pestle size suitable for nuclei conservation. Lysates were transferred to low-binding protein tubes and centrifuged for 5 minutes at 2500xg and 4°C. The resulting pellets were washed twice with the appropriate restriction enzyme ice-cold buffer and resuspended in 360 µl of ice-cold buffer. Undigested input samples were saved to verify digestion efficiency. Following this, SDS was added to 0.1%, and nuclei were incubated at 65°C for 10 minutes. After immediate incubation on ice, Triton X-100 was added to 1% for SDS quenching. At this point, 200 U of the restriction enzyme (BtgI, NEB) was added, and samples were incubated overnight at 37°C with interval shaking.

Digested samples were incubated for 20 minutes at 65°C to inactivate residual enzymatic activity. 10 µl aliquots were saved as digestion controls for comparison with undigested samples. The 3C protocol was optimized to achieve >85% digestion efficiency in all experiments. Ligation was performed by adding 720 µl of the ligation mix (400 U T4 DNA ligase, 1.7x T4 DNA ligase buffer, 1.7% Triton X-100, 200 µg/ml molecular-grade BSA) and incubating the samples for 4 hours at 16°C with interval shaking. To reverse the crosslinking, Proteinase K was added to a final concentration of 0.5 mg/ml and samples were incubated for 2 hours at 65°C. A second round of Proteinase K addition was performed before an overnight incubation at 65°C. Samples were then purified by phenol-chloroform and ethanol precipitation in 300 mM AcNa and 70% EtOH. A second purification step with G-50 Sephadex beads was performed to clean the samples from salts and contaminants before RNAse A treatment (30 min, 37 °C, 40 µg/ml). dsDNA was quantified using Qubit, and 50 ng of DNA was loaded per qPCR plate well. Internal primers amplifying undigested regions were used for loading adjustments.

For normalization purposes, a bacterial artificial chromosome (BAC) clone (RP11-884L15) containing regions of interest was digested and ligated following the same procedure. This allowed normalization of the data to the intrinsic frequency of ligation for each pair of regions. The selected BAC clone covers all genomic regions studied in this work. BACs were purified from 500 ml overnight cultures. 20 µg of DNA were digested, ligated and ligation frequencies determined as previously described for test samples. Interaction frequencies were expressed as the ratio between the ligation products obtained in experiments and the ligation products obtained from the BAC.

### Biotinylated-Psolaren Immunofluorescence

Biotinylated-psoralen (bPsoralen) incorporation was employed to evaluate (-) DNA supercoiling, following the methodology outlined in ^43^. Briefly, cells were cultured on polylysine pre-treated glass slides and exposed to 75 µM bPsoralen (EZ-Link Psoralen-PEG3-Biotin, ThermoFisher) along with 0.01% digitonin (Sigma, D141) for 20 minutes at room temperature in dark conditions. After 10 minutes of UV crosslinking at 360 nm, slides were fixed in 4% PFA-PBS for 15 minutes at room temperature. Then, slides were blocked in 5% BSA-PBS before incubating with primary antibodies (1:2500 α-ERα mouse antibody and 1:1000 α-biotin) for 30 minutes. Coverslips were then washed three times in 0.1% Tween20-PBS and incubated with secondary antibodies for 30 minutes. DAPI counterstaining and ProLong mounting (ThermoFisher) were performed for image acquisition using a LEICA confocal microscope SP5. Quantification of bPsoralen was restricted to nuclei (DAPI) or intranuclear regions exhibiting a high density of ERα signal (HDR). As negative controls, cells were pretreated with triptolide (10 µM, 5 h).

### Immunoprecipitation

For co-immunoprecipitation (IP) experiments, cells were seeded in 15 cm plates at 80% confluence. Following treatment, culture plates were washed with cold PBS, and cells were scraped and centrifuged at (300xg, 5 min, 4°C). The resulting cell pellets were resuspended in 600 µl of ice-cold lysis buffer (20 mM Tris-HCl pH 8, 0.5% Triton-X100, 2 mM MgCl_2_, 2.5 kU benzonase, Protease Inhibitor Cocktail, and PMSF) and incubated on ice for 30 minutes. Simultaneously, 4 µg of IP antibody were incubated with Dynabeads (12.5 µl Protein-A and 12.5 µl Protein-G) at room temperature for 40 minutes in IP buffer (20 mM Tris-HCl pH 7.5, 200 mM NaCl, 0.5 mM EDTA, Protease Inhibitor Cocktail, 1 mM DTT). After lysis, cell extracts were centrifuged (13000 rpm, 5 min, 4°C), and 0.5 mg of total protein from the supernatant was diluted in IP buffer (to reduce Triton-X100 concentration below 0.2%) and incubated in rotation with antibody-bound Dynabeads (2 h, 4°C). Upon incubation, beads were washed three times with IP buffer and boiled for 10 minutes in 2x Laemmli Buffer before conducting western blot analysis.

### Western blotting

Protein extracts were prepared by boiling cells in lysis buffer (125 mM Tris-HCl pH 6.8, 20% (w/v) glycerol, 4% SDS) for 10 minutes. Lysates were then mixed with Laemmli buffer and subjected to an additional boiling step. Approximately 15-20 µg of protein were loaded onto either 10% in-house acrylamide gels or 4-20% Mini-PROTEAN Tris-Glycine Precast Protein Gels (BioRad). Samples were then transferred to an Immobilon-FL Transfer PVDF Membrane (Millipore) for 2 hours at 4°C and 70V.

To block nonspecific binding, membranes were incubated in 5% BSA-TBS for 1 hour at room temperature. This was followed by an overnight incubation with primary antibodies at 4°C in 5% BSA-TBS with 0.1% Tween20. Afterwards, membranes were washed three times in TBS with 0.1% Tween20. And incubated with the corresponding IRDye-conjugated secondary antibody in 5% BSA-TBS with 0.1% Tween20. After three washes with buffer, membranes were dried and subjected to analysis using the Odyssey CLx system and ImageStudio Odyssey CLx software (LI-COR BIOSCIENCES, Lincoln, NE), following the manufacturer’s protocols.

### TOP2B Plasmid Relaxation assay

The negative supercoiled plasmid relaxation assay was conducted using the Inspiralis kit (HTRB201). Recombinant human TOP2B (2.8 μM) was incubated with 0.5 μl of supercoiled pBR322 plasmid (1 μg/μl) for the specified duration, following standard procedures. Human recombinant ZATT was added to the reaction at a final concentration of 100 nM. To stop reactions, 30 μl of STOP solution (containing 40% (w/v) sucrose, 100 mM Tris-HCl pH 8, 1 mM EDTA, 0.5 mg/ml Bromophenol Blue) was added. Subsequently, the reactions were purified and cleaned by adding an equal volume of chloroform/isoamyl alcohol (24:1, v:v). After vortexing and centrifugation, the aqueous solution was loaded onto a 1% (w/v) agarose gel for 2 hours at 85V. Subsequently, gels were stained with ethidium bromide (1 μg/ml in water) and visualized using a transilluminator.

### *In vitro* SUMOylation assay

*In-vitro* TOP2B SUMOylation assay reactions (20 μl) were conducted in a buffer containing 20 mM HEPES pH 7.6, 50 mM NaCl, 0.05% Tween 20, 1 mM DTT, 200 μM ATP, and 4 mM MgCl_2_. The reaction mixture consisted of 30 nM TOP2B (Inspiralis), 3 μM SUMO2 (S294-31H-200, Signalchem), 1 μM UBC9, 100 nM SUMO E1 mix (SAE1/UBA2, BML-UW9330-0025, ENZO), and 10 nM ZATT. Reactions were incubated at 30°C for 90 minutes and assessed by Western Blot using anti-TOP2B antibodies. For the relaxation assays, 3 μl of the SUMOylation reaction mixture was used, following the previously described procedure.

### Chromatin immunoprecipitation (ChIP)

To prepare chromatin, 10 cm culture plates at 80% confluency were crosslinked using 1% formaldehyde (Sigma) in cell medium for 10 minutes at 37°C. The crosslinking reaction was quenched with 125 mM glycine (Sigma). Subsequently, culture plates were washed twice with ice-cold PBS, and cells were collected in ice-cold PBS supplemented with 1x Complete Protease Inhibitor Cocktail (Roche) and 1 mM PMSF using cell scrapers. Cells were then centrifuged (300xg, 5 min, 4°C) and lysed in two steps. In the first step, cells were incubated for 10 minutes in 1 ml of Lysis Buffer A (5 mM Pipes pH 8, 85 mM KCl, 0.5% NP40, 1 mM PMSF, 1x Complete Protease Inhibitor Cocktail) for nuclei isolation. Next, Lysis Buffer B (50 mM Tris HCl pH 8.1, 1% SDS, 10 mM EDTA, 1 mM PMSF, 1x Complete Protease Inhibitor Cocktail) was added to the pelleted nuclei (4000 rpm, 5 min) for nuclear lysis. Sonication was performed using Bioruptor (Diagenode, UCD-200) at high intensity, with 25 cycles (30 seconds sonication followed by a 30-second pause). Chromatin was then clarified by centrifugation (17,000xg, 10 min, 4°C).

For assessing chromatin fragmentation, 5% of the chromatin volume was incubated in Lysis Buffer B with 0.25 mg/ml Proteinase K (AppliChem) for 16 hours at 65°C to reverse the crosslinking. The DNA was subsequently purified using phenol:chloroform extraction and ethanol precipitation (70%) and quantified by Nanodrop 1000 (ThermoFisher). Fragment size distributions were analyzed by agarose gel electrophoresis to obtain a smear ranging between 300-600 bp.

For IP, 30 µg of chromatin and the corresponding amount of antibody (according to STAR Methods) were incubated overnight in IP buffer (20 mM TrisHCl pH 8, 150 mM NaCl, 0.1% SDS, 1% TritonX-100, 2 mM EDTA, 1x Complete protease inhibitor cocktail, 1 mM PMSF) at 4°C. Subsequently, 25 µl of BSA pre-blocked protein A and protein G Dynabeads (ThermoFisher) were added and incubated with chromatin (4h, 4°C). The lysis buffer was diluted at least 5 times in IP buffer to reduce SDS concentration to less than 0.2%. Beads were then sequentially washed with rotation using IP buffer, IP buffer with increased salt (500 mM NaCl), LiCl buffer (20 mM TrisHCl pH 8, 0.25 M LiCl, 1% NP40, 1% NaDoc, 1 mM EDTA), and TE buffer. To elute the DNA, beads were incubated at 50°C in 100 µl elution buffer (1% SDS, 100 mM NaHCO_3_). Crosslinking was reversed by incubating samples with 200 mM NaCl and 100 µg of Proteinase K (ThermoFisher) (65 °C, overnight). The DNA was purified using Qiagen PCR purification columns (QUIAGEN, 28106) after treatment with RNAseA (0.5 mg/ml). Finally, samples were subjected to qPCR using the indicated primers (according to STAR Methods). Data is presented as a percentage of input (mean +/- SEM), with individual dots representing independent biological replicates.

### Chromatin immunoprecipitation and sequencing (ChIPseq)

ChIPseq protocol followed the previously described ChIP method with small variations. For spike-in normalization, mouse chromatin from MEFs was added to the samples before immunoprecipitation (IP) at a 1:20 ratio. Upon LiCl washing step, chromatin bound Dynabeads were subjected to the ChIPmentation protocol as described in Schmidl et al. (2015). In brief, beads were washed twice with 10 mM TrisHCl pH 8. Then, samples were transferred to new tubes for incubation with Tn5 enzyme in buffer (50 mM TrisHCl pH 8, 50% (v/v) DMF, 25 mM MgCl2) at 37°C for 10 minutes to incorporate adaptors into chromatin fragments. The Tn5 enzyme used was provided by the Proteomic Service of CABD (Centro Andaluz de Biología del Desarrollo). Following tagmentation, samples were immediately placed on ice, and washed again with LiCl buffer and TE buffer before transferring to new tubes.

Next-generation sequencing (NGS) libraries were prepared after quantifying and assessing the quality of the DNA fragments by qPCR. Illumina-indexed primers, which partially align with the Tn5-introduced adapters, were used in qPCR to validate Tn5 reactions and quantify DNA amount. Optimal Cqs (quantification cycle) were determined for the actual library preparation based on the qPCR results, and the libraries were then amplified for N cycles, where N corresponds to the starting point of the exponential curve amplified in the qPCR. qPCR reactions were performed using an in-house SyberGreen master mix under the same conditions as the library PCR. Libraries were generated using NEBNext High-Fidelity Polymerase (NEB, M0541). Subsequently, size selection of DNA fragments ranging from 300 to 500 bp was performed using Sera-Mag Select Beads (GE Healthcare, 29343052). A 0.7x volume of beads mixture was used to remove fragments larger than 500 bp, followed by the addition of an additional 0.15x volume to isolate fragments larger than 200 bp. Primers were added at a final concentration of 0.2 µM, and PCR amplification was performed according to the following protocol: 1x (72°C, 5 min), 1x (98°C, 30 s), Nx (98°C, 10 s - 63°C, 30 s - 72°C, 30 s), 1x (72°C, 5 min). NGS libraries were sequenced at the Genomic Unit of CABIMER (Seville, CSIC) using a NextSeq 500 HIGH-Output flowcell in a single-end configuration with 75 bp read length. Prior to sequencing, libraries were quantified using Qubit and analyzed for size profiles using Bioanalyzer.

### *In vivo* complex of enzyme assay (ICE)

Cells, treated and untreated with Etoposide (VP16, Sigma, E1383) or Camptothecin (CPT, C9911), were lysed immediately in a denaturing cell lysis solution consisting of 1% (v/v) N-Lauroylsarcosine sodium salt (Sigma-Aldrich, L7414) and 1x Complete Protease Inhibitor Cocktail. Cell plates were scraped, and lysates were collected into polypropylene tubes. Samples were homogenized using a latex-free syringe with a 25G needle, and genomic DNA precipitation was performed through a CsCl density gradient.

For DNA precipitation, CsCl (Applichem-Panreac, A1098) was added to lysates to achieve a final concentration of 0.67 g/ml. Lysates were ultracentrifuged (57000 rpm, 20 h, 25°C) using 3.3 ml polyallomer Optiseal tubes (Beckman Coulter) in a TLN100 rotor (Beckman Coulter). After centrifugation, DNA pellets were washed with 1 ml of 70% EtOH and resuspended in molecular-grade H_2_O containing 1x Complete Protease Inhibitor Cocktail. Samples were incubated at 4°C for 30 minutes to ensure complete rehydration of DNA, followed by a 5-minute heating step at 65°C in a water bath. DNA quantification was performed using a Nanodrop 1000. To remove nucleic acids, samples were incubated with benzonase (NEB) for 30 minutes on ice. Then, DNA and RNA-free samples were diluted in TE buffer before transferring them to a nitrocellulose membrane (Odyssey Nitrocellulose Membrane, LI-COR Biosciences) using the Bio-Dot SF microfiltration apparatus (Bio-Rad). Immunodetection was performed following the same protocol as described for Western Blot experiments.

### ICE-IP and ICEseq

To map topoisomerase cleavage complexes, material isolated by ICE assay was subjected to IP. In this process, 40 µg of ICE isolated DNA was digested using dsFragmentase (NEB). Samples were incubated with 10 µl of the enzyme in 1x dsFragmentase buffer and 10 mM MgCl_2_ according to the standard protocol (30 min, 37 °C). Enzymatic reactions were stopped by adding 50 mM EDTA. The size of the resulting DNA fragments was assessed using agarose gel electrophoresis to obtain a DNA smear ranging from 200 to 500 bp. Subsequently, samples were diluted in IP buffer, and the immunoprecipitation was performed as in ChIP protocol with minor modifications. Incubations with 2 µg of anti-topoisomerase antibodies were carried out in the presence of 0.5 mg/ml molecular-grade BSA. DNA was eluted from beads in 100 µl elution buffer (30°C, 30 min) and treated with 100 µg of Proteinase K (37°C, 2 h). Samples were then purified using Sera-Mag SpeedBeads magnetic beads (Fisher) in a 1:1 ratio following standard procedures before qPCR analysis.

The ICEseq protocol included additional variations. Firstly, isolated topoisomerase cleavage complexes bound to antibody-coated dynabeads were subjected to the ChIPmentation protocol, as described for ChIPseq. Secondly, immunoprecipitated and purified DNA was subjected to a second step of Tn5 tagmentation before library preparation.

### High-throughput sequencing analysis

Sequenced reads obtained from NGS were demultiplexed and quality filtered using FastQC and subsequently aligned against either human (hg19) or mouse (mm9) genome utilizing Bowtie 1.2 ^64^. The option "-m 1" was used to retain only uniquely mapped reads. "Rsubread" package was used to align ICEseq reads with specific parameters (type = 1, TH1 = 2).

To correct technical biases and global differences, spike-in normalization was applied using mouse chromatin as control in human cell line experiments, following the methodology described in ^65^. Then, signal tracks were obtained using bamCoverage (deepTools) without normalization and with a bin size of 50 bp. To measure the enrichment ratio in mouse and human signal tracks, the ratio between the IP and input samples was calculated. This approach helps to avoid local biases and background noise, providing a more accurate representation of the occupancy ratio (OR). Thus, OR was obtained by dividing the enrichment ratio in mouse and human reads upon IP. For the genome tracks, this OR was used to normalize signal tracks using bamCoverage with RPGC normalization (50 bp bin size and 300 bp read extension). For ChIPseq quantification, the OR obtained from spike-in normalization was applied to RPGC generated using bedCoverage (Samtools) at sites of interest.

ICEseq signal tracks were generated with RPKM normalization, using a bin size of 50 bp and a read extension of 300 bp. For quantification of ICEseq data, differential binding analysis was performed through the integration of R libraries, including ’csaw’, ’edgeR’, ’rtracklayer’, and ’statmod’. Bam files were used as input with the specific parameters: window width = 150, spacing = 50 and minq = 40. ‘normFactors’, ‘asDGEList’, and ‘estimateDisp’ functions were used for normalization and differential binding analysis using CPM values. Subsequently, the ‘glmQLFit’ and ‘glmLRT’ functions were used for comparative tests. When only one experimental dataset was available, the dispersion was set to 0.05 and a comparative test was performed using the ‘glmFit’ algorithm.

For correlation analysis, the corresponding regions were systematically sorted and grouped into non-overlapping sets containing of 5-10 regions. This strategy aimed to mitigate the potential impact of outliers on the analysis. Subsequently, values were correlated using the Spearman method in R.

Visualization of signal tracks was performed using the UCSC Genome Browser, with mean values and a smoothing window between 6-12 pixels. Signal profiles and heatmaps were generated using ^66^. For plot profiles, signal values were represented by the mean in bins of 100 bp. We used Iterative Grubbs’ method (alpha = 0.2) to remove outliers when quantifying sequencing data at specific loci.

MACS2 was used for calling broad peaks from ChIPseq experiments ^67^. For annotation, called peaks were extended 1.5 kb bidirectionally. Regulatory regions, namely enhancers, promoters, and insulators, were defined as follows. For promoters, the whole set of transcripts associated with Ensembl-annotated genes were considered. Then, promoters were defined as ± 1 kb from the TSSs. Enhancers were defined as H3K27ac peaks not overlapping with a promoter, and insulators as CTCF peaks not overlapping with promoters and enhancers.

To obtain E2-responsive enhancers, annotated enhancers from the GROseq dataset [GSE43836] were used ^3^. The enhancers were called according to the following criteria. Transcripts larger than 5 kb were identified and annotated in the RefSeq, ENSEMBL, and UCSC Known Gene databases to classify already known annotated transcripts. Transcripts on the same strand that overlapped with the existing annotation of protein-coding genes, tRNA, rRNA, snoRNA, or miRNA were discarded. The enhancer coordinates annotated in the hg18 genome assembly were converted to the hg19 assembly using the UCSC LiftOver tool. Transcripts that ran antisense to gene annotations and transcripts that overlapped with the promoter regions of annotated transcripts (window +/- 500 bp) were also rejected. Protein-coding genes were obtained from annotated transcripts in the ENSEMBL database (hg19). Induced and non-induced enhancers were classified based on RNA POLII ChIPseq data from four independent experiments (45 min, 10 nM E2). This classification was performed using bedCoverage analysis with Samtools (v.1.6) and R. Induced genes and enhancers were defined as those exhibiting a log2 fold change (log2FC) greater than 1 and a p-value less than 0.05 (based on a t-test with the greater alternative hypothesis). E2 non-responsive genes and enhancers were randomly selected to match the same number as induced transcripts, considering transcripts with log2FC values between -0.1 and 0.1, p-values greater than 0.05, and RNA POLII values greater than 1^e-3^. To identify ERα-mediated contacts co-localizing with enhancers, the ChIAPET dataset was intersected with ER peaks annotated at enhancers (peaks overlapped with H3K27ac peaks but not Ensembl-annotated genes).

## QUANTIFICATION AND STATISTICAL ANALYSIS

Data presented in this study originates from independent biological replicates, individual dots in each graph signifies each replicate. Detailed information regarding data collection, quantification and statistical methods are provided within the corresponding sections in method details. Statistical analyses were performed using GraphPad Prism software v.9 (GraphPad, San Diego, CA, USA) and R (v.3.2.0).

## Notes

### Competing Interest Statement

The authors have declared no competing interest.

